# Visualizing the translation landscape in human cells at high resolution

**DOI:** 10.1101/2024.07.02.601723

**Authors:** Wei Zheng, Yuekang Zhang, Jimin Wang, Shuhui Wang, Pengxin Chai, Elizabeth J Bailey, Wangbiao Guo, Swapnil C Devarkar, Shenping Wu, Jianfeng Lin, Kai Zhang, Jun Liu, Ivan B Lomakin, Yong Xiong

**Affiliations:** Department of Molecular Biophysics and Biochemistry, Yale University, New Haven, CT 06511, USA; Microbial Sciences Institute, Yale University, West Haven, CT 06516, USA; Department of Microbial Pathogenesis, Yale University, New Haven, CT 06536, USA; Department of Pharmacology, Yale University, West Haven, CT 06516, USA; Department of Dermatology, Yale University, New Haven, CT 06520, USA

## Abstract

Obtaining comprehensive structural descriptions of macromolecules within their natural cellular context holds immense potential for understanding fundamental biology and improving health. Here, we present the landscape of protein synthesis inside human cells in unprecedented detail obtained using an approach which combines automated cryo-focused ion beam (FIB) milling and *in situ* single-particle cryo-electron microscopy (cryo-EM). With this *in situ* cryo-EM approach we resolved a 2.19 Å consensus structure of the human 80S ribosome and unveiled its 21 distinct functional states, nearly all higher than 3 Å resolution. In contrast to *in vitro* studies, we identified protein factors, including SERBP1, EDF1 and NAC/3, not enriched on purified ribosomes. Most strikingly, we observed that SERBP1 binds to the ribosome in almost all translating and non-translating states to bridge the 60S and 40S ribosomal subunits. These newly observed binding sites suggest that SERBP1 may serve an important regulatory role in translation. We also uncovered a detailed interface between adjacent translating ribosomes which can form the helical polysome structure. Finally, we resolved high-resolution structures from cells treated with homoharringtonine and cycloheximide, and identified numerous polyamines bound to the ribosome, including a spermidine that interacts with cycloheximide bound at the E site of the ribosome, underscoring the importance of high-resolution *in situ* studies *in* the complex native environment. Collectively, our work represents a significant advancement in detailed structural studies within cellular contexts.

## Introduction

Structure determination of biological macromolecules at high resolution within their native cellular environment is an enticing pursuit. Advances in cryo-electron tomography (cryo-ET) have enabled steady progress toward this goal^1,2,3^. Recent applications of cryo-ET to thin cellular samples (∼100-200 nm thick), such as small bacteria or cellular lamellae milled by a focused ion beam (FIB)^4,5,6^, have reached 3-4 Å resolution for large macromolecular complexes such as ribosomes^7,8,9^. While increasingly powerful and efficient, cryo-ET is still time-intensive, low-throughput, and limited in resolution. *In situ* single-particle cryo-EM (Fig. 1a), has emerged as a promising method to interrogate large cellular complexes inside the cell, especially complexes that are transiently stable, heterogenous, or otherwise hard to purify and reconstitute *in vitro*^10,11^. However, the full potential of the in situ single-particle cryo-EM for new biological discovery had not yet been demonstrated.

**Fig. 1:**
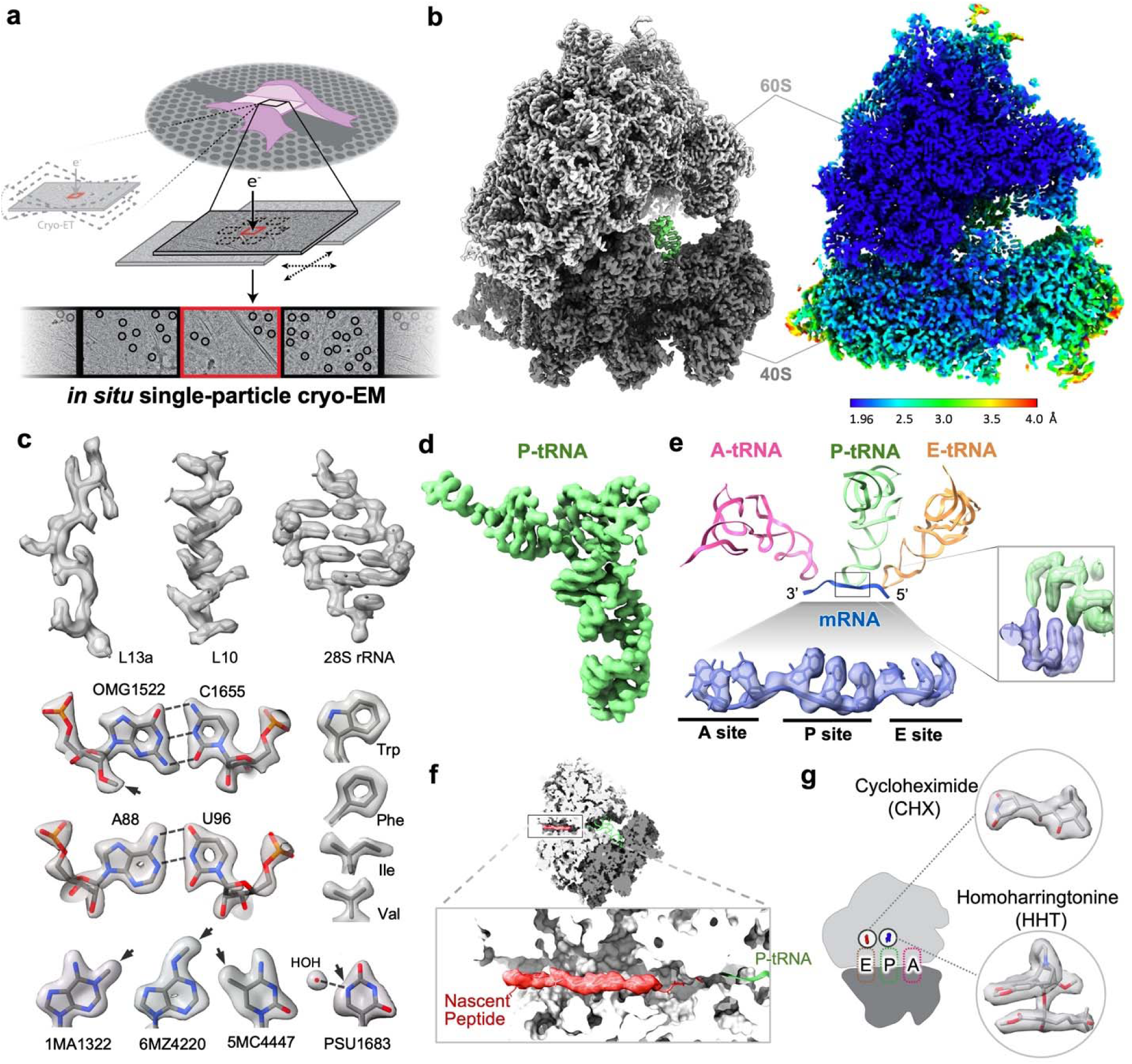
High-resolution *in situ* structure of the human ribosome. **a,** Schematic of the *in situ* single-particle cryo-EM pipeline. **b,** Consensus map of human 80S ribosome at 2.19 Å resolution (left) and the local resolution map (right). **c,** Features of the 80S ribosome structure. The cryo-EM map is shown as a semitransparent surface with the fitted model in stick representation. rRNA modifications are marked with arrows. **d,** Well-resolved cryo-EM map of the P site tRNA (P-tRNA). **e,** Cryo-EM density of mRNA (a polyU model, blue sticks) in A, P, and E sites of the 40S subunit (bottom) and its interaction with the P-tRNA (right inset). The well-resolved density for mRNA is an average of all translating mRNAs. A schematic of the interactions is shown on the top. **f,** Density for the nascent peptide (red) in the 60S subunit. The ribosome is shown as a gray surface clipped at the nascent peptide exit tunnel. **g,** Cryo-EM densities of homoharringtonine (HHT) and cycloheximide (CHX) (shown by sticks) resolved in cells treated with both drugs.

The ribosome is a prime example of a compositionally heterogeneous and conformationally dynamic macromolecular machine. The eukaryotic ribosome comprises the small and large subunits (40S and 60S subunits, respectively), and translates mRNA into protein through a sequence of initiation, elongation, termination, and recycling steps^12,13,14,15^. During elongation, synthesis of the nascent peptide requires intricate concerted processes, including small subunit rotation, cofactor coordination, tRNA translocation, and movement of mobile structural elements^13,15^. Extensive *in vitro* studies of purified ribosomes using X-ray crystallography and cryo-EM have provided much of our current understanding of translation^12,15^. However, ribosomes are often purified in stable states such as hibernating states^16^. Affinity purification and *in vitro* assembly have been used to study some specific ribosomal complexes^17,18,19,20,21,22^. Many dynamic states have been captured by *in vitro* studies using ribosomal inhibitors, providing access to an extensive cycle of translation elongation^9,22,23,24,25^. Recent *in situ* cryo-ET studies have revealed aspects of ribosome dynamics inside living cells, including drug interactions and their impact on translation^7,8,9^, albeit with limited details of the functional states and the biological insight gained.

Using *in situ* single-particle cryo-EM, we achieved a detailed 2.19 Å resolution consensus structure of the human 80S ribosome, identifying 21 unique conformations with bound tRNAs and cofactors, and identified 21 unique 80S ribosome conformations with bound tRNAs and cofactors, resolving 20 conformations to below 3 Å resolution and one to 3.26 Å. This provides an unprecedented view of the translation cycle. We identified ribosome-bound protein factors, such as endothelial differentiation related factor 1 (EDF1) and nascent polypeptide-associated complex /3 (NAC/3), that were not previously known to be enriched on purified ribosomes. Most strikingly, we identified new structural regions of SERBP1 and found its N terminal region binds at all ribosome states and C terminal region binds at all ribosome states except states containing A/P-P/E tRNAs. The SERBP1 can connect the 60S and 40S ribosomal subunits, suggesting its roles in translation regulation. In addition, we resolved several higher-order assemblies of translating ribosomes in details, including a detailed di-ribosome interface that is consistent with helical polysomes previously observed at low resolution^26,27–29^. Additionally, we analyzed structures from cells treated with homoharringtonine and cycloheximide, and identified numerous polyamines bound to translating ribosomes, including one that interacts with cycloheximide bound in the ribosomal E site. Our results underscore the potential of the *in situ* single-particle cryo-EM approach for high-resolution structural studies of biological complexes in their native environment.

### High resolution *in situ* human 80S ribosome structure

To investigate the structure of the human ribosome at inside the cell, we used cryo-FIB milling to section 193 lamellae from human cells (HEK293A), from which we collected 16,636 micrographs using single-particle cryo-EM data collection procedure. Particle picking and initial orientation determination were first performed by template matching with a 60S ribosomal subunit reference map^30^, followed by further search with the refined 80S map, using the program GisSPA^23,31^ and cisTEM^32^ (Fig. 1a; Extended Data Table 1). 3D classification and refinement using the single-particle pipeline in CryoSPARC^33^ yielded a 2.19 Å consensus cryo-EM map of the 80S ribosome, with the highest local resolution below 2 Å (Fig. 1b, Extended Data Fig. 1c). The density for ribosomal proteins and ribosomal RNA (rRNA) were resolved at a level that has not been previously achieved *in situ* (Fig. 1c), including numerous rRNA modifications consistent with previously identified *in vitro* (Fig. 1c)^30,34–36^.

The exceptional quality of the cryo-EM map enabled us to discern factors, including inhibitors and drugs, that were bound to ribosomes. Focused classification on the tRNA, mRNA, and nascent peptide allowed us to resolve densities for these features in 80S ribosomes (Fig. 1d-f). The path of the mRNA through the A (aminoacyl), P (peptidyl) and E (exit) tRNA-binding sites is clearly visible, including well-resolved densities for nucleotide bases that are averaged from all translating mRNAs in the cell (Fig. 1d, e). We also applied this pipeline to cells treated with homoharringtonine (HHT) and cycloheximide (CHX) and obtained a consensus structure (2.42 Å resolution) of the 80S ribosome with the two drugs bound. The quality of the map allowed us to unambiguously model HHT and CHX (Fig. 1g). The level of information obtained here far surpasses the previous *in situ* ribosome structures and underscores the effectiveness of combining single-particle cryo-EM with FIB-milling to obtain high-resolution structures of biomolecular complexes in their native surroundings in the cell for drug discovery.

### Translation dynamics in human cells

This large dataset allowed us to capture a comprehensive depiction of translation dynamics *in situ*. The 80S ribosome passes through an array of compositionally and conformationally distinct states during translation elongation to select the cognate aminoacyl-tRNAs and add the amino acids to the growing nascent peptide chain^12,13,14,15^. Using focused classification, we identified 21 distinct ribosome states, all but one resolved to under 3 Å resolution (Fig. 2a, b, Extended Data Fig. 2, Extended Data Table 1). Of these, 14 states represent actively translating ribosomes, as indicated by clear density for the mRNA, tRNAs, and protein factors, and are consistent with those described in the literature (Fig. 2a, Extended Data Fig. 3a)^7,8,9,29,37^. The most prevalent states are the eEF1A-A/T-P-E, eEF1A-A/T-P-Z, and eEF1A-A/T-P states (states 2a, 2b, 2c, respectively; Fig. 2a) which together comprise 30.3% of all 80S ribosomes. The mRNA is disordered in these states, suggesting that these are codon sampling states where the mRNA has not yet formed stable hydrogen bonds with the cognate tRNA (Extended Data Fig. 3a). The higher overall occupancy of these eEF1A-A/T states is consistent with the notion that proofreading is a rate-limiting step in translation elongation^38^. We also resolved the high-resolution structures of the ribosome with eEF1A in a post GTP hydrolysis state (states 2a’, 2b’, 2c’), where the mRNA is structured and interacts with the A/T-tRNA (Extended Data Fig. 4b). This is accompanied by a large rotation of the A/T-tRNA, which now interacts with the helix 89 (H89) of the 28S rRNA, as opposed to H43 and H44 before rotation (Extended Data Fig. 4a).

**Fig. 2:**
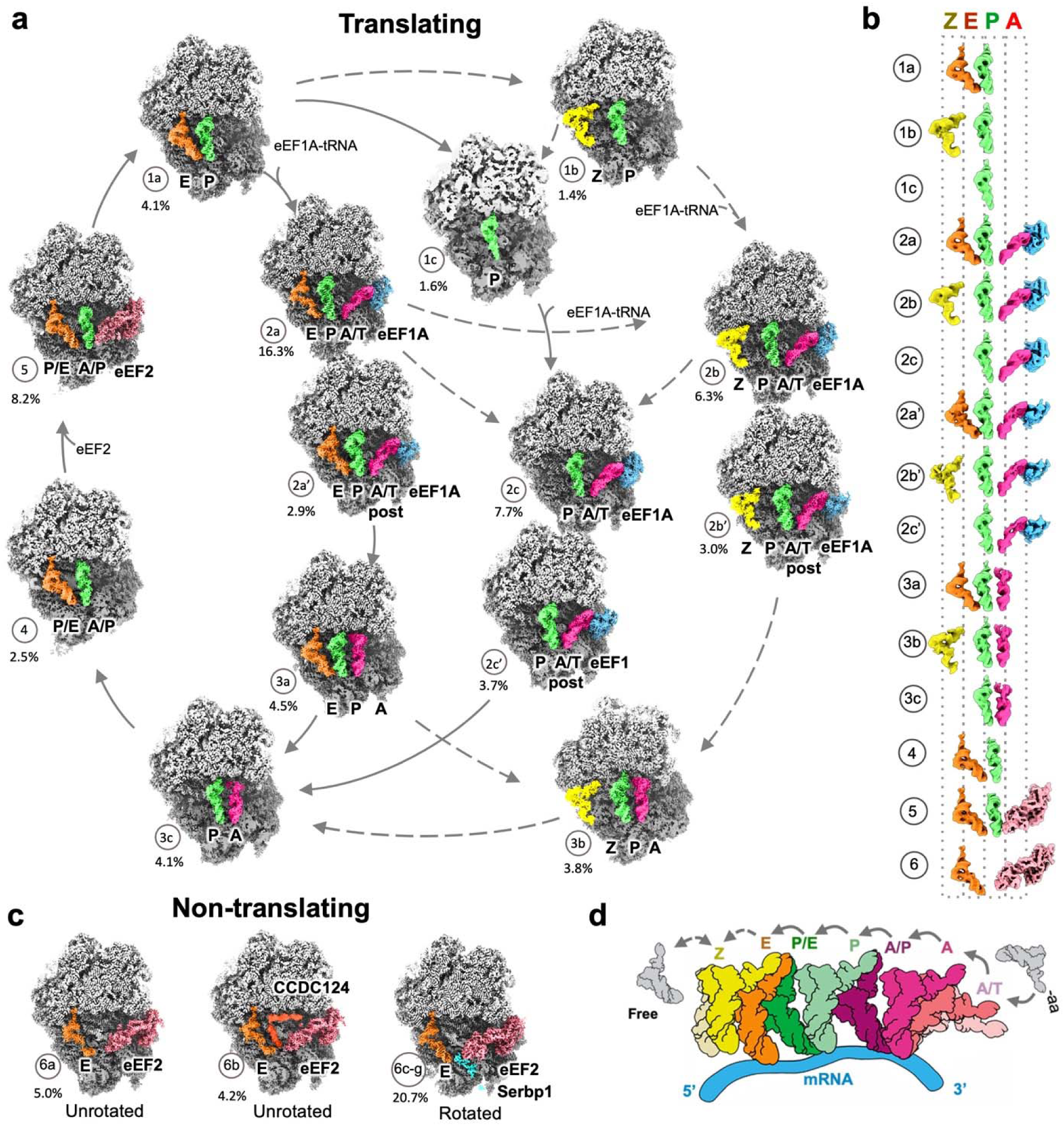
Translation dynamic during elongation. **a**, 14 classes of human 80S ribosome in the translation elongation cycle, clipped to show tRNAs and factors bound. All but one were resolved to a resolution higher than 3 Å, except for state 1c at 3.26 Å resolution. Solid arrows indicate pathways supported by literature. Dashed arrows indicate possible pathways. The percentages indicate the proportion of particles in each class. The 60S and 40S subunits are shown in light and dark gray respectively. The tRNAs are colored by position. **b**, Comparison of the positions of tRNAs and translation elongation factors in each class, colored as in 2a. **c**, 7 classes of hibernating states, with the SERBP1 (cyan)-constraining states (6c-g) grouped together. **d**. Schematics for the movement of a tRNA through the ribosome during translation elongation.

Additionally, we observed 7 non-translating ribosome states with bound E site tRNA and eEF2. Among these, two states feature an unrotated 40S subunit (states 6a, 6b; Fig. 2c), with the translation reactivation factor CCDC124 bound in one of the states (6b). The remaining five states are associated with SERBP1 and exhibit varying degrees of 40S subunit body rotation and head swivel, shedding light on the action of SERBP1 (described below). (states 6c-g; Fig. 2c, Extended Data Fig. 5d, f).

Our data also provides insights into the functional roles of the recently discovered noncanonical Z site in the ribosome^9,29,39^. We resolved four distinct states with a tRNA bound at the Z site (Z-tRNA, states 1b (P-Z), 2b (eEF1A-A/T-P-Z), 2b’ (Post-eEF1A-A/T-P-Z) and 3b (A-P-Z); Fig. 2a, b). In agreement with a previous report^39^, we observed that the Z-tRNA interacts with both the 60S and 40S ribosomal subunits and its position is incompatible with the presence of another tRNA in the E site (Extended Data Fig. 4c, d). The acceptor arm of the Z-tRNA interacts with eL42 and H68 of the 28S rRNA, while the anticodon loop is held by eS25 (Extended Data Fig. 4c). Two of the three Z-tRNA states (1b and 2b) have similar conformations, but the anticodon loop of the tRNA in the third state (3b, Z’) and eS25 shift towards the solvent together by ∼3.8 Å. The T-loop/elbow of the Z-tRNA is bound to the backbone of the L1 stalk, similar to how the E-tRNA contacts the L1 stalk, and the stalk moves with the tRNA toward the Z site (Extended Data Fig. 4d). Therefore, it is possible that the L1 stalk facilitates the displacement of tRNA from the E site through a rigid-body motion, guiding it toward the solvent via the Z site. Future studies will need to interrogate this possibility.

### Newly identified structure and function of SERBP1

Our *in situ* cryo-EM maps enabled us to observe new structural features and binding modes of SERBP1 (Fig. 3). Previously, two small central regions of SERBP1 were observed to bind hibernating ribosomes^16,40^ and the density of the remainder of the protein was not resolved on purified ribosome. Here, we resolved both the N- and C-terminal regions of SERBP1 anchored to the 60S and 40S subunits, respectively. The N-terminal regions of SERBP1 are bound to all 80S ribosomes (Fig. 3a) and free 60S subunits (Fig. 3b). The C-terminal regions bind to all ribosomes except those in A/P-P/E and A/P-P/E-eEF2 states. Both binding sites are supported by clear density and independent AlphaFold-2^41,42^ predictions (Fig. 3c). Based on the new structures, we propose a model in which SERBP1 is always bound to the 60S ribosome subunits through its N-terminal regions, while the C-terminal regions bind the 40S except in the rotated PRE states (A/P-P/E and A/P-P/E-eEF2). In this manner the central region of SERBP1 is poised to occupy the mRNA channel and rapidly shuttle the ribosome between translating and hibernating states (Fig. 3d). It has been reported previously that SERBP1 binds free 40S subunits^43^. If SERBP1 engages both the free 60S and 40S subunits simultaneously, it could act as a flexible tether between the two (Fig. 3d).

**Fig. 3:**
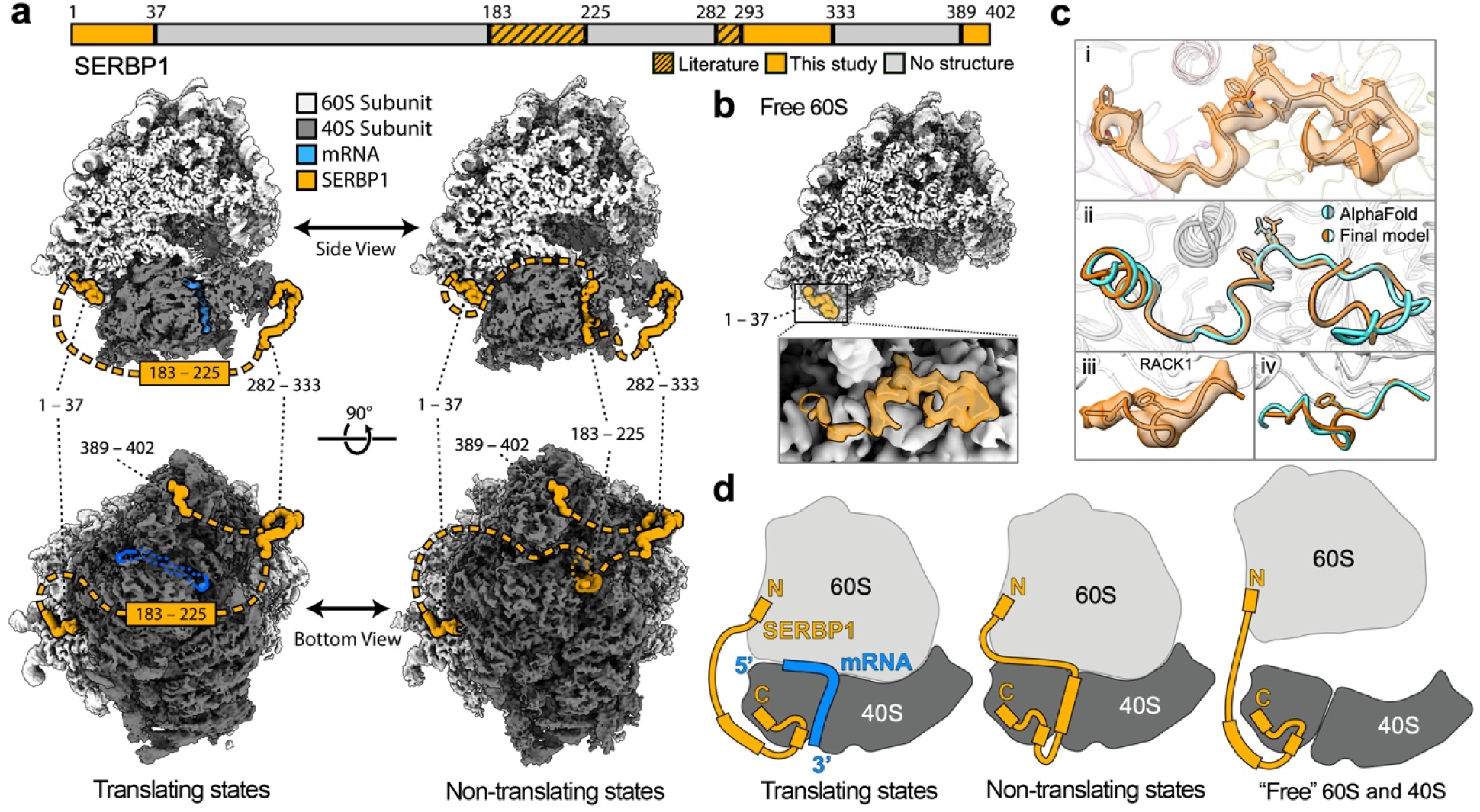
Newly identified binding sites of SERBP1 and functional implications. **a,** Structure of SERBP1 (orange) in translating and non-translating ribosome states. The central region of SERBP1 was resolved in hibernating states in the mRNA entry channel as in previous studies, while the N-terminal and C-terminal regions were identified in this study. **b,** The N-terminal region of SERBP1 also binds to the free 60S subunit. **c,** The atomic models of the N-terminal (i, ii) and C-terminal (iii, iv) regions of SERBP1 fit well in the densities and match closely to the AlphaFold-2 predictions (cyan). **d,** Cartoon models for the roles of SERBP1 in translation, in which it links ribosomal subunits in all states, potentially even when they are separated as individual subunits.

Analysis of hibernating states with different 40S subunit body rotation and 40S head swivel shows no density for the SERBP1 central region in the unrotated state (state 6a; Fig. 3c), weak density in a state with highest 40S subunit head swivel and lowest 40S subunit body rotation (state 6c; Extended Data Fig. 6a), and strong density in higher 40S subunit body rotation (state 6d-g; Fig. 3c). Based on these states, we propose that the 40S subunit head swivel opens the mRNA entry channel to allow the central region of SERBP1 to slot into the channel. Then, the 40S subunit body rotates to close around the mRNA channel and stabilize the binding of the SERBP1 central region (Extended Data Fig. 6a, Extended Data Movie. 1).

The N-terminal region of SERBP1 is localized near the ribosome collision interface. To examine whether SERBP1 binding is affected by ribosomal collision, we used focus classification on the space beyond the mRNA exit channel of the 40S subunit and observed multiple structures of colliding ribosomes (Extended Data Fig. 6b). We found that when both the stalled and collided ribosomes are in the A/P-P/E states, the rRNA H79 helix of the leading ribosome exhibits a conformational change near the binding site of the N-terminal region of SERBP1. We observed that the density for the SERBP1 N-terminal region reduces on the leading ribosome in these interactions (Extended Data Fig. 6b). This suggests that SERBP1 may be involved in sensing and regulating ribosome collision.

### Identification of new interaction interface between adjacent translating ribosomes

The polysome was identified in the mid 1960^44^ and it often form helical assemblies^45,46^. Short segments with two or three colliding ribosomes, known as disomes and trisomes respectively, were also frequently observed^47–49^. 3D structures of helical polysomes in both prokaryotic and eukaryotic species were studied by cryo-ET at moderate resolutions^26–29^. The disome interface was resolved to high resolution using purified short polysomes, although this interface cannot form the helical structure^50^ (Fig. 4e). Here, we analyzed the polysome densities of different ribosome states. All translating states show some polysome density at the mRNA entry and exit positions (Extended Data Fig. 7A). Interestingly, the A/P-P/E states show strong polysome densities at the mRNA entry position, indicating that these states are enriched in adjacent ribosomes. Additional classification of di-ribosomal particles resolved two main classes of di-ribosomes with different interaction interfaces at resolutions between 3 to 4 Å (Fig. 4a, b, Extended Data Fig. 8). One class (class2, “top-top” interaction) is similar to previously published disome interfaces and has mainly 40S-40S subunit interactions^7,9,50^ (Fig. 4a, b). The other class (class1, “top-back” interaction) is distinguished by a large rotation of the leading ribosome relative to the trailing ribosome (Fig. 4a, b, c). This interface is mainly formed by 60S-40S subunit interactions and is similar to the helical arrangement revealed by previous cryo-ET studies at low resolution (∼25 Å) ^28,29^. When the cryo-EM data were analyzed using a large box size, we indeed observed helical polysome density (Fig. 4d) and fitting the class1 di-ribosome structures can form the helical polysome structure consistent with previous analysis^29^ (Fig. 4e). The leading ribosomes with the class2 interaction consist of both unrotated and rotated translation states which should be actively translational form for polysome (Extended Data Fig. 8), similar to that recently visualized in *E. coli* polysome^51^. By contrast, the compact helical polysomes consist of only specific A/P-P/E states for both leading and trailing ribosomes, which can only be inactive for translation. It has been reported that inhibition of protein synthesis and virus infected cell can induce helical polysomes^46,52^. We observed fewer than 0.5% of the total ribosomes in helical polysomes, potentially pausing translation to adapt to the local cellular environment.

**Fig. 4:**
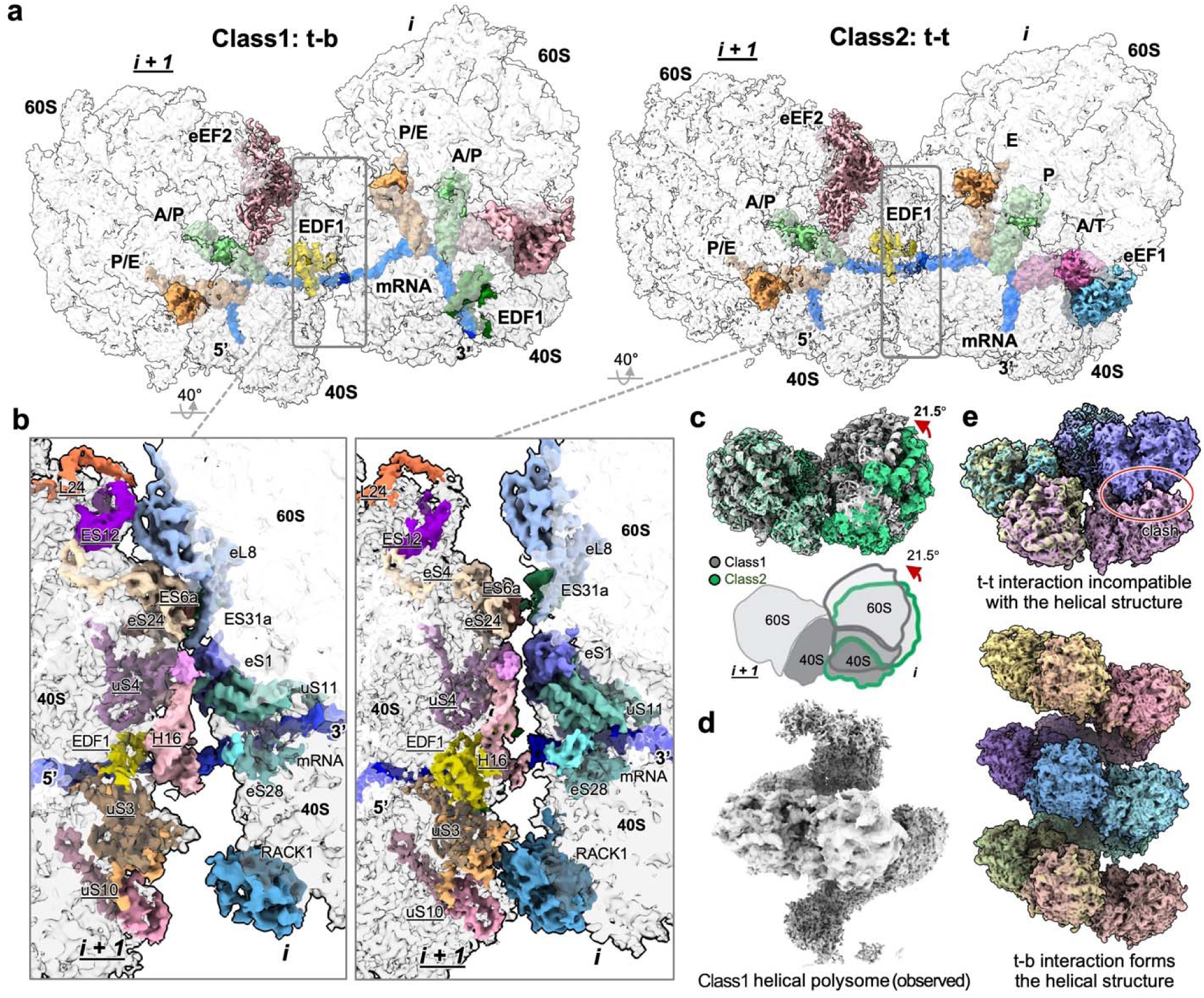
Identification of new interaction interface of di-ribosomes. **a,** Structures of two types of di-ribosomes. *i* indicates the leading ribosome, *<u>i+1</u>* indicates the trailing ribosome. Class1 is a top-back (t-b) interaction and Class2 is a top-top (t-t) interaction. **b,** Interaction interfaces of the two types of di-ribosomes. The t-b class has mainly inter-ribosomal 60S-40S subunit interactions, whereas the t-t class has mainly 40S-40S subunit interactions. **c,** Comparison of the two classes of di-ribosomes. **d,** Helical class1 ribosome density appears when data processing was done with a large box. **e**, The t-t interaction is incompatible with the helical structure shown in d, whereas the t-b interaction can form this high-order helical ribosome structure.

### Ribosome-binding cofactors identified *in situ*

Our cryo-EM maps allow us to identify many factors bound to ribosomes *in situ*, including EDF1, NAC/3, ErbB3-binding protein 1 (EBP1), and the eukaryotic initiation factor 6 (eIF6) (Fig. 5, Extended Data Fig. 9). EDF1 is known to be involved in resolving ribosomal collisions^20^. In our cryo-EM maps EDF1 is distinctly visible on approximately half of the ribosomes that associate with a preceding ribosome (Fig. 4b). Surprisingly, in our EDF1-bound ribosome structures, helix 16 (h16) of the 18S rRNA displays a different conformation compared to the EDF1-bound structure from purified samples^20^ (Fig. 5b). It appears to be “clamped” by the N- and C-terminal regions of EDF1 in a way that has not be observed before (Fig. 5b). These changes may be functionally important and highlight the importance of conducting structural biology *in situ*.

**Fig. 5:**
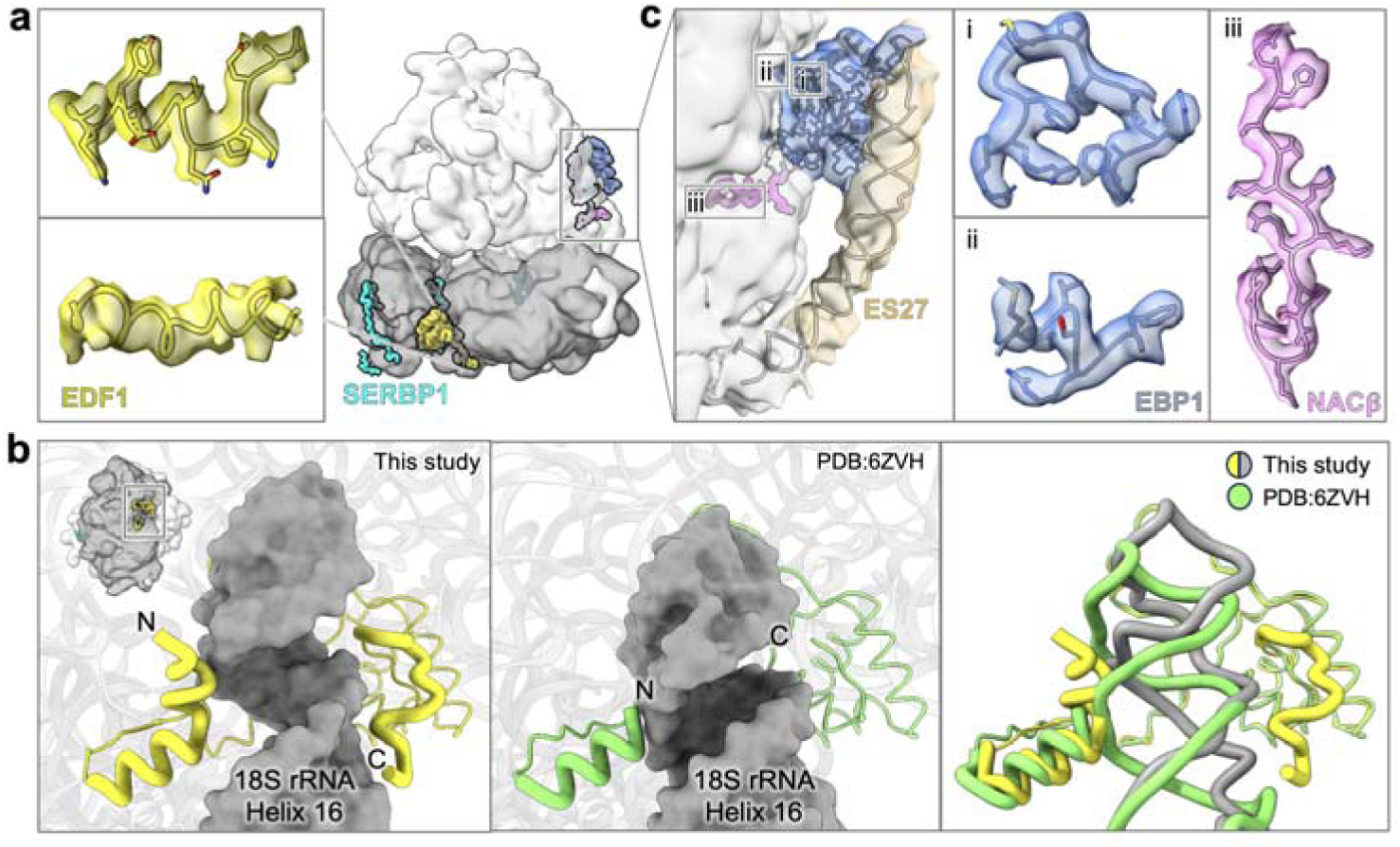
Ribosome binding factors in cells. **a,** Ribosome binding factors EDF1, SERBP1, EBP1, and NAC/3 were identified on ribosomes inside the cell. Representative cryo-EM maps (surface) and structural models (cartoon and sticks) are shown. **b,** Conformations of h16 of 18S rRNA and EDF1 observed *in situ* and *in vitro*. **c,** NAC/3 and EBP1 binding sites.

NAC/3 is involved in nascent peptide processing and trafficking but has only been observed in reconstituted samples *in vitr*o^21^. Our maps show that the N-terminal anchor domain of NAC/3 is enriched on all translating ribosome states with occupancies of ∼50% (Fig. 5c, Extended Data Fig. 9a). The C-terminal globular domain of NAC/3, which we did not observe, has been reported to compete with the binding of EBP1, a protein associated with ribosomal translational regulation at the nascent peptide exit channel^53^. We observed that EBP1 and NACβ N-terminal occupations exhibit modest inverse correlation (Fig. 5c, Extended Data Fig. 9a). In hibernating 80S ribosomes and free 60S subunits, the occupancies of EBP1 increase, whereas those of NACβ drop. This correlation might be important for the nascent peptide exit and translation regulation. Furthermore, unlike the previous cryo-ET study which did not observe eIF6 on 60S subunits^9^, our 60S subunit structure showed clear density for eIF6 bound to the inter-subunit interface (∼60% occupancy), poised to prevent 40S ribosomal subunit joining (Extended Data Fig. 9b).

### Native ligand and drug binding in the human ribosome

We further demonstrated that the *in situ* single-particle cryo-EM approach can be efficiently applied to the study of the detailed effects of pharmaceutical agents in cells. We treated 293A cells with both the ribosome inhibitor cycloheximide (CHX), which competes with tRNA for binding at the E site^24^, and the anticancer drug homoharringtonine (HHT), which prevents peptide bond formation after aminoacyl-tRNA binding at the A site^24,54^. With our approach, we obtained a 2.42 Å 80S ribosome consensus map with clear densities for both compounds, allowing for atomic-model building for the drug-ribosome interactions (Fig. 6).

**Fig. 6:**
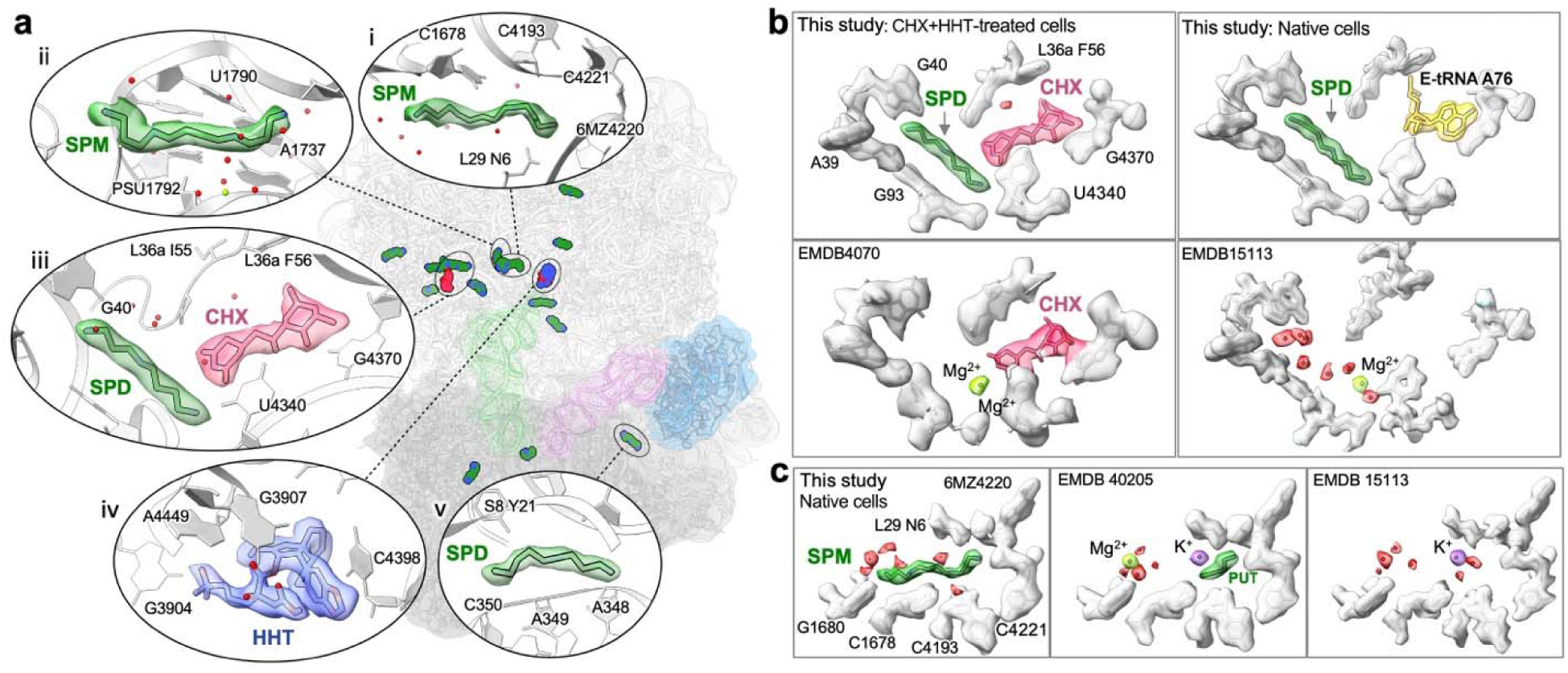
Native ligand and drug binding in the human ribosome. **A,** 14 polyamines were identified in native human ribosomes and those in CHX- and HHT-treated cells. Cryo-EM densities (surface) and stick models for representative polyamines and drugs (HHT and CHX) are shown with selected surrounding ribosome residues (cartoon). **b,** Comparison of the spermidine molecule contacting CHX with compositions of the same binding pocket observed in *in vitro* studies. **c,** Comparison of a spermine molecule observed *in situ* with compositions of the same binding pocket observed in *in vitro* studies.

Our *in situ* cryo-EM maps allowed us to provide the first description of native ligand binding on the human ribosome, in both native cells and the CHX- and HHT-treated cells. We identified 14 polyamine molecules, including spermine and spermidine, bound to the ribosome (Fig. 6, Extended Data Table 2). In previous *in vitro* structures of purified ribosomes, the identity of ligands at these sites was influenced by buffers used during purification^30,38,55^, and thus are often different from what we observed *in situ*. Of particular note, a spermidine molecule contacts CHX and alters the chemical environment of its binding pocket (Fig. 6a-iii, 6b). This exemplifies the importance of *in situ* ligand characterization for drug design and screening, as well as for understanding the role of ligands in in translation regulation.

By employing focused classification and refinement, we resolved seven ribosome states in the CHX+HHT-treated cells at 3 - 4 Å resolutions, each with well-resolved drug densities (Extended Data Fig. 10, 11). Six of the observed states corresponded to translating ribosomes (states 1b, 1c, 2b, 2c, 3b, 3c; Extended Data Fig. 11) and accounted for ∼58% of 80S ribosomes, while a single hibernating state (state 6h, Extended Data Fig. 11) accounted for the remaining ∼42% of 80S. As anticipated, no state harbored a tRNA in the E or P/E sites, consistent with the inhibition of tRNA translocation from the P to the P/E site by these two inhibitors. Our observations from the drug-treated cells also provided insight into the mechanistic role of the Z-tRNA. Nearly 50% of translating ribosomes (28.7% of all ribosomes) in the drug-treated cells contained Z-tRNA. This is higher than the proportion of Z-tRNA-containing states in the untreated cells (27.3% of translating ribosomes, 22.2% of all ribosomes) (Extended Data Fig. 11). Partially anticipated, this outcome is attributed to the CHX- and HHT-mediated depletion of the E site tRNA, which competes with tRNA binding at the Z site (Extended Data Fig. 4d). Importantly, the CHX and HHT treatment potently inhibited translation, and resulted in substantial presence of tRNA binding at the Z site. The elevated Z-tRNA level did not rescue translation inhibition at the E site, indicating that it is not an independent pathway of tRNA release bypassing the E site requirement. Instead, it is consistent with the Z site being a subsequent step in translation that occurs after E site tRNA binding events, potentially also involving an equilibrium with the rebinding of deacylated free tRNA (Fig. 2d).

## Discussion

The combination of automated cryo-FIB milling and single-particle cryo-EM methods holds the potential to transform the study of biomolecules within their natural cellular contexts. As a complement to cryo-ET, it offers a faster and more efficient approach to obtain well-resolved structures. In our large-scale study, we highlight the power of combining the template matching (GisSPA and 2DTM) and the conventional single-particle cryo-EM workflow to facilitate *in situ* structural biology investigations with both high throughput and high resolution, achieving comparable resolution to those achieved with *in vitro* purified samples. More importantly, we capture of macromolecular states within the cells that are unattainable *in vitro*, thereby providing new biological insights. This approach can be effectively employed for detailed investigation of inhibitors within cells, holding profound implications for new drug development.

With these extensive structures of the *in situ* translation landscape, we have resolved factors that are not often found in purified ribosome, including NAC/3, and EDF1. Strikingly, we identify new binding modes of SERBP1, showing that it binds both the 40S and 60S subunits, perhaps to allow for swift switches between translating and non-translating ribosome states. Bridging the free 60S and 40S subunits, it may also allow for rapid assembly of 80S ribosomes during translation initiation. Even more provocatively, our observation that all individual 60S subunits are bound by SERBP1 indicates that the ribosomal subunits may be constantly tethered, a possibility that needs to be interrogated by future studies. With our large dataset, we provide a comprehensive translation elongation cycle and a robust distribution of ribosome states inside the cell. In addition, we have determined detailed structures of the native ribosome-ribosome interface during translation, which has proven challenging for other methods including cryo-ET. We also elucidated the detailed interaction between adjacent ribosomes that form helical polysome structures, which is likely a minor, inactive form of translation that pauses to adapt to the cellular environment.

As structural biology expands its exploration of the cellular interior, it will be interesting to see how observations will differ across different organisms and cell lines. A striking feature of *in situ* data is that the raw datasets contain the full repertoire of cellular factors and, thus, can be reanalyzed for entirely different macromolecular targets. This versatility opens exciting possibilities for structural biology to uncover the mechanisms of various cellular processes as they occur in the native cell. The continued development of efficient, high-resolution *in situ* structural biology tools will undoubtedly accelerate this exploration.

## Author contributions

Y.X. supervised the project. W.Z. prepared the sample and performed FIB milling with the help of J.F.L., J.L. and W.G.. W.Z., S.P.W. and Y.X collected the data. W.Z., P.C., S.H.W., Y.Z., K.Z. and Y.X. processed the data. W.Z., J.W., Y.Z., S.H.W., S.C.D., E.J.B, I.L. and Y.X., interpreted the data. W.Z., J.W., Y.Z. built and refined the atomic models. W.Z., Y.Z., S.H.W., E.J.B., S.C.D., and Y.X. prepared the figures. W.Z., E.J.B., Y.Z., I.L and J.W. wrote the manuscript with input from all authors. J.W., Y.Z., E.J.B., I.L., S.H.W., W.Z. and Y.X. edited the manuscript.

## Acknowledgements

The project is funded by Yale discretionary funds to Y.X. The Aquilos Cryo-FIB is funded by R01AI152421 (J.L.). We thank Yale CryoEM Resource for maintaining the Krios Cryo-TEM used for data collection. We thank Yale Center for Research Computing for support with computing resources. We thank colleagues at the Department of Molecular Biophysics and Biochemistry at Yale University for suggestions and discussions on this study.

## Data availability

The atomic models of the untreated consensus, the consensus focused refined on 40S head, the consensus with EBP1, 60S, 60S with eIF6, state 1a, 1b, 1c, 2a, 2b, 2c, 2a’, 2b’, 2c’, 3a, 3b, 3c, 4, 5, 6a, 6b, 6c, 6d, 6e, 6f, 6g, the HHT and CHX treated consensus, consensus with EBP1, 60S, 60S with eIF6, state 1b, 1c, 2b, 2c, 3b, 3c, 6h, the t-t and t-b disome consensus composite have been deposited to the Protein Data Bank (PDB) under the accession codes 9AZC, 9AZM, 8UJB, 8UJC, 8UJD, 8UIZ, 8UIY, 8UJ9, 8UJR,8UJQ, 8UJ8, 9AZS, 9B0O, 9B0N, 8UJ5, 8UJ4, 8UJ3, 9AZN, 9B0W, 9B11, 9B0J, 9B0P, 9B0R, 9B0F, 9B0G, 9B0H, 9B0V, 8UIL, 8UIM, 8UK0, 8UJP, 8UJO, 8UJN, 8UJK, 8UJJ, 8UKB, 9B0S, 9B0Q. The globally and locally refined cryo-EM maps have been deposited to the Electron Microscopy Data Bank (EMDB) under accession codes EMD-44014, EMD-44016, EMD-42317, EMD-42318, EMD-42319, EMD-42306, EMD-42305, EMD-42316, EMD-42326, EMD-42325, EMD-42315, EMD-44021, EMD-44048, EMD-44047, EMD-42312, EMD-42311, EMD-42310, EMD-44017, EMD-44060, EMD-44063, EMD-44043, EMD-44049, EMD-44051, EMD-44039, EMD-44040, EMD-44041, EMD-44059, EMD-42298, EMD-42299, EMD-42339, EMD-42324, EMD-42323, EMD-42322, EMD-42321, EMD-42320, EMD-42351, EMD-44052, EMD-44057, EMD-44056, EMD-44050, EMD-44054, EMD-44053. Also see Extended Data Table 1. Other data are available from corresponding authors upon reasonable request.

## Supplementary

**Extended Data Fig. 1:**
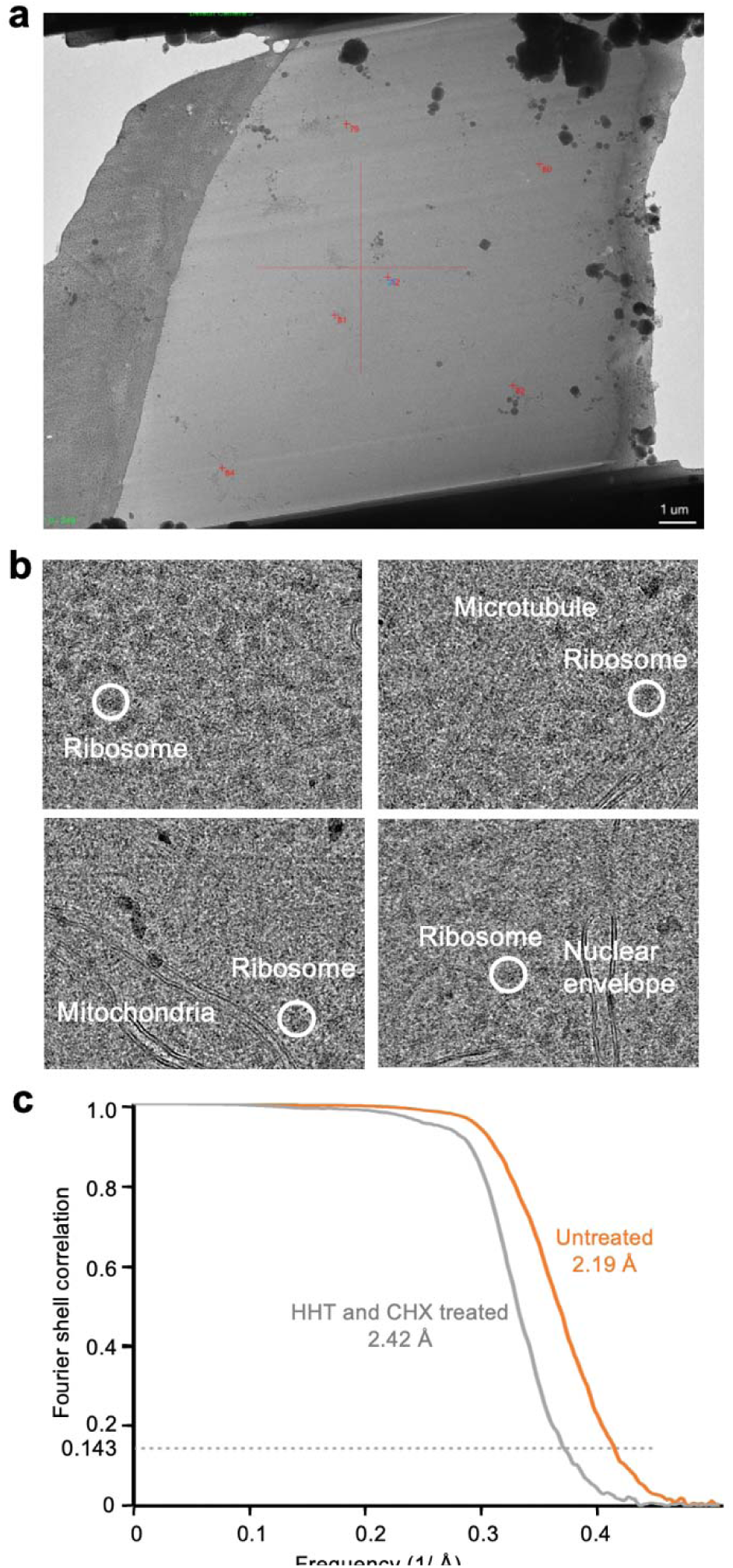
*In situ* single particle cryo-EM. **a,** A representative transmission electron micrograph of FIB-milled lamellae. **b,** Typical micrographs collected by *in situ* single particle cryo-EM. Prominent features such as ribosomes, microtubules, mitochondria, and nuclear envelope can be visibly identified. **c,** The Fourier shell correlation (FSC) curves of the consensus 80S reconstruction with data form untreated 293A cells.

**Extended Data Fig. 2:**
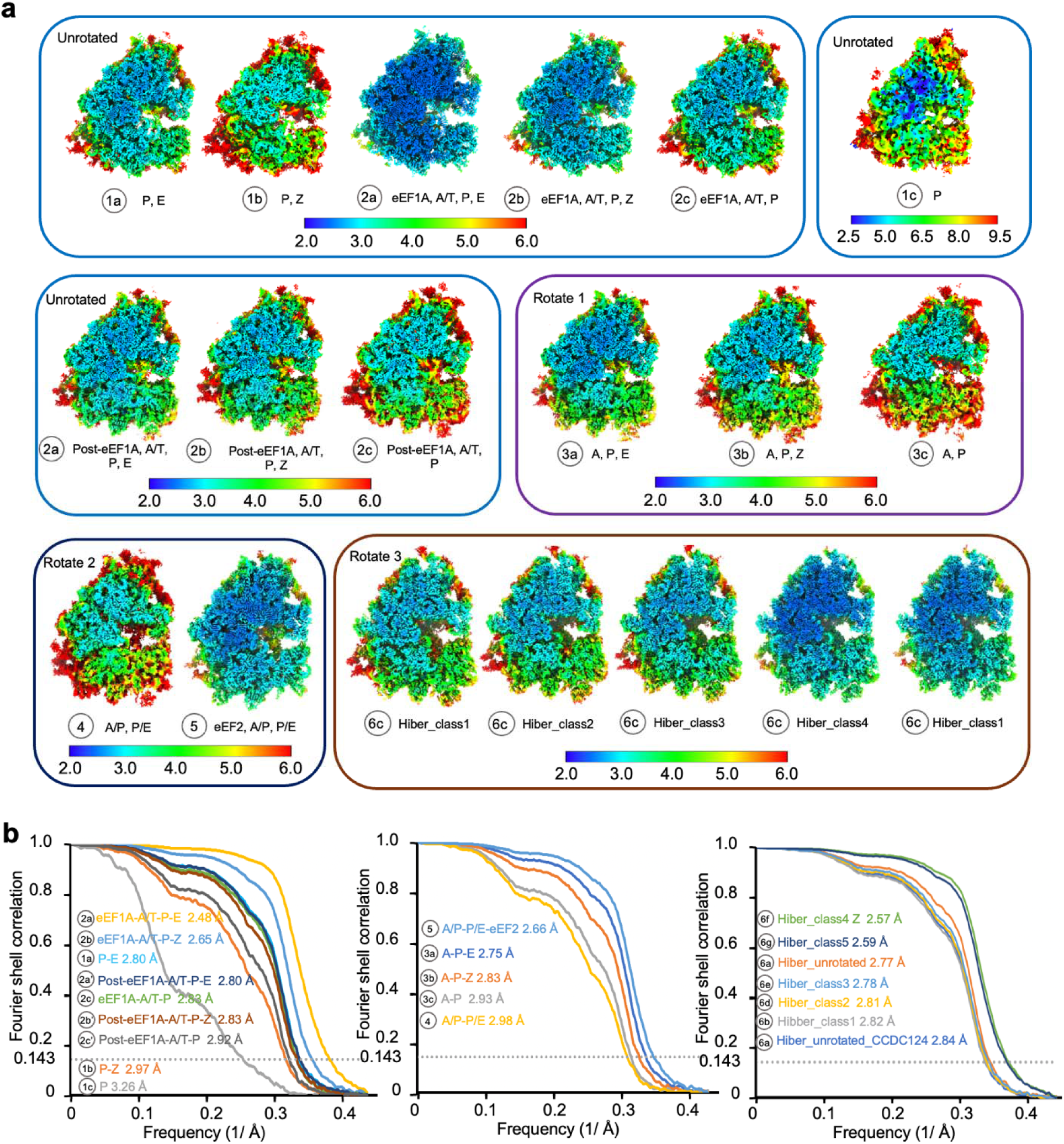
Local resolution maps and FSC of ribosome states. **a,** Local resolution maps of ribosome states in the untreated cells. **b,** FSC curves of the untreated ribosome states using the 0.143 criterion.

**Extended Data Fig. 3:**
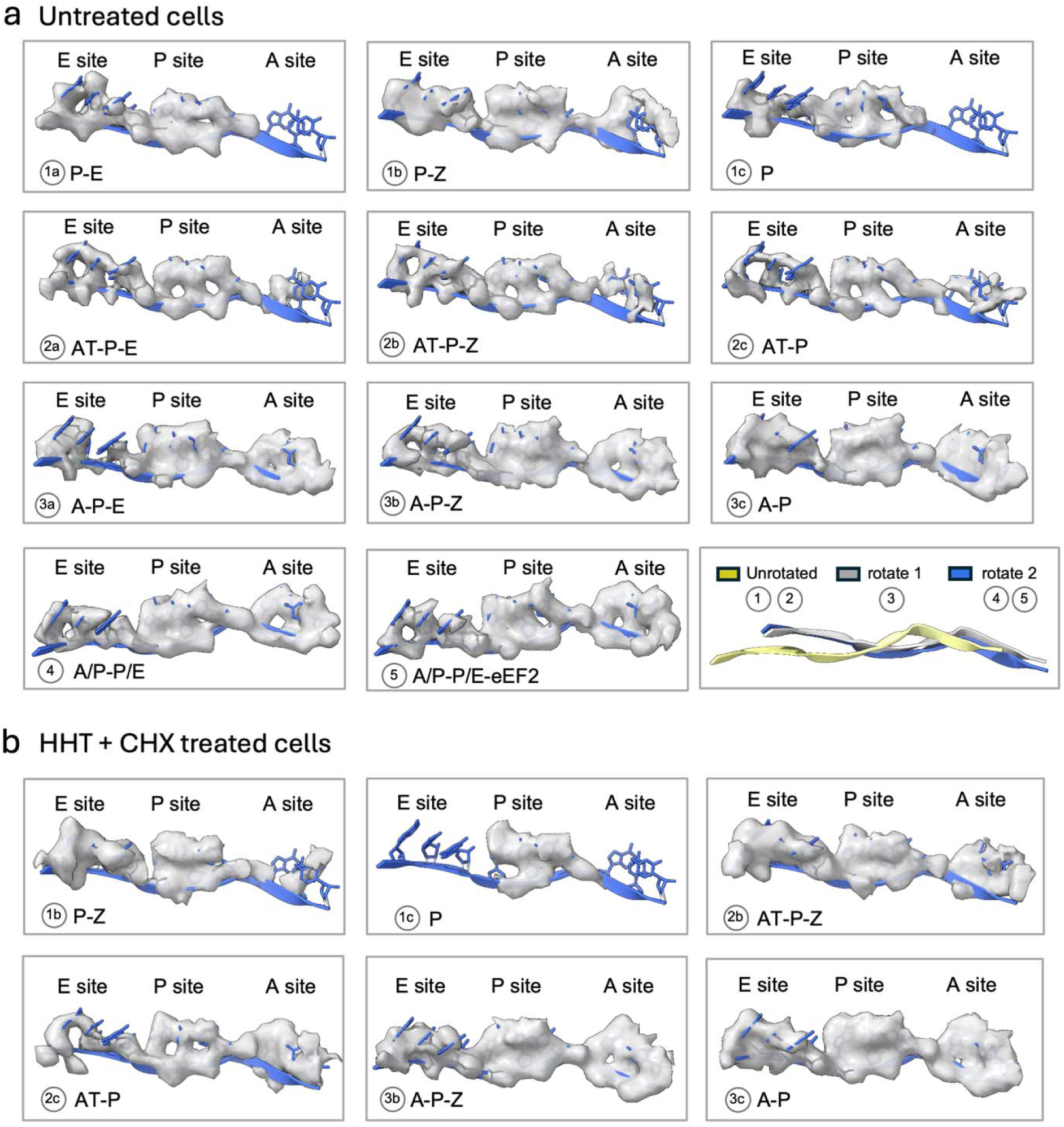
mRNA densities at the A, P, and E sites in different ribosome states. **a,** mRNA densities (white density) at A, P, E site in different ribosome states from the untreated dataset. The last panel shows the relative positions of mRNA in different rotation states. **b,** mRNA densities at A, P, and E sites in different ribosome states from the HHT+CHX-treated dataset. The same contour level is used for all panels.

**Extended Data Fig. 4:**
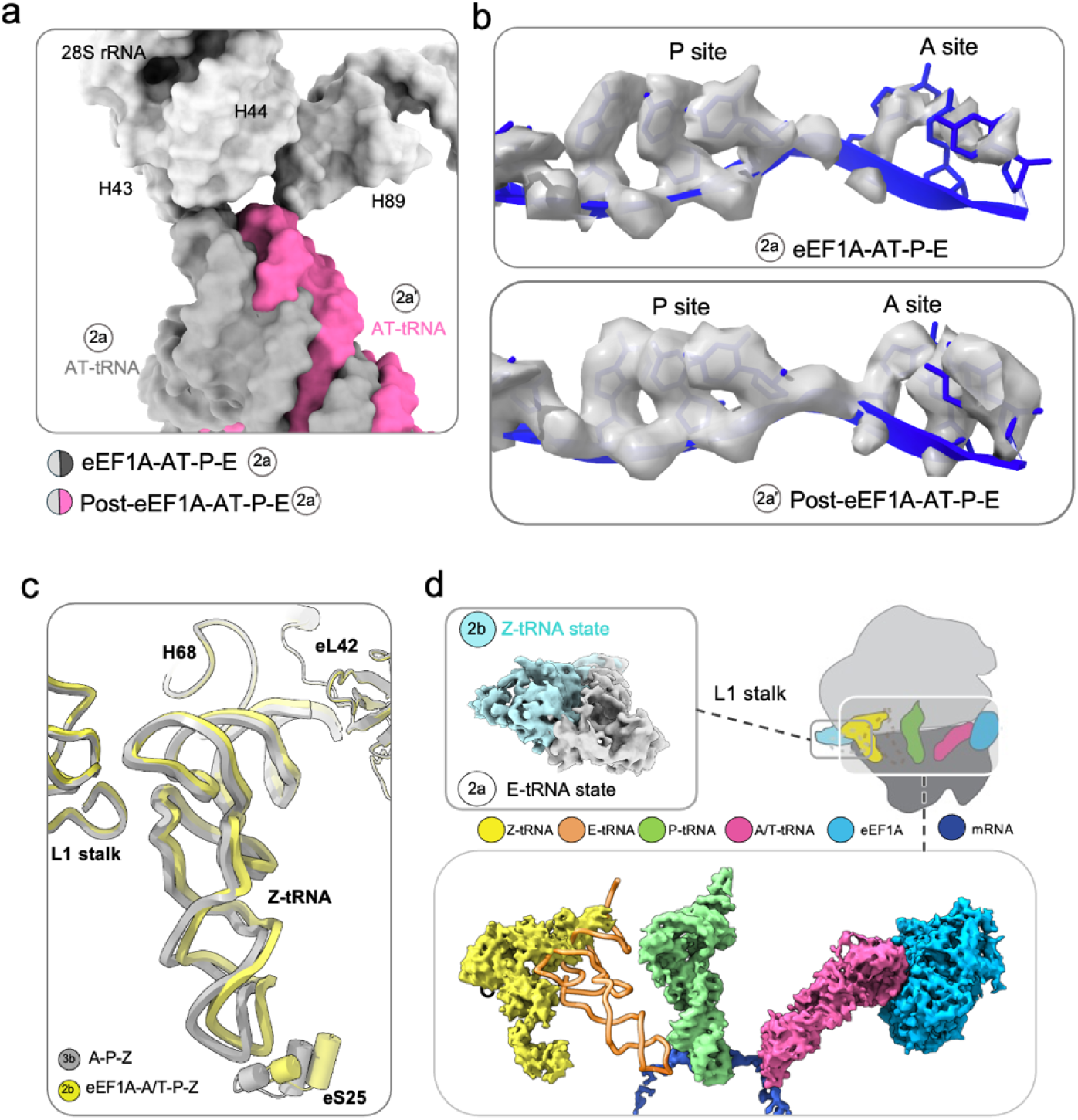
The AT-tRNA and Z-tRNA in active translation ribosomes. **a**, AT-tRNA movement between the codon-sampling (2a) and the post GTP hydrolysis (2a’) states shows different interactions of tRNA with the 60S ribosomal subunit. b, The mRNA in the post GTP hydrolysis state (bottom) is better resolved comparing with the codon-sampling states (top). **c**, The Z-tRNA is held by the L1 stalk, eS25, H68, and eL42. The anticodon loop of the Z-tRNA and eS25 are displaced by ∼3.8 Å in the A-P-Z (3b) state (grey) relative to its position in the eEF1-A/T-P-Z (2b) state (yellow). **d**, A comparison between the eEF1-A/T-P-Z state (2b) and the eEF1-A/T-P-E state (2a). Top inset: cryo-EM maps of the L1 stalk (surfaces) showing it moves towards the solvent in state 2b (cyan) relative to its position in state 2a (gray). Bottom inset: Unlike the E-tRNA (orange ribbon), the Z-tRNA (yellow density) does not contact the mRNA (dark blue density). The cryo-EM map of the tRNAs and eEF1A, along with the mRNA in the eEF1-A/T-P-Z (2b) state is shown in surface representation. An E-tRNA model (orange ribbon) is shown for comparison.

**Extended Data Fig. 5:**
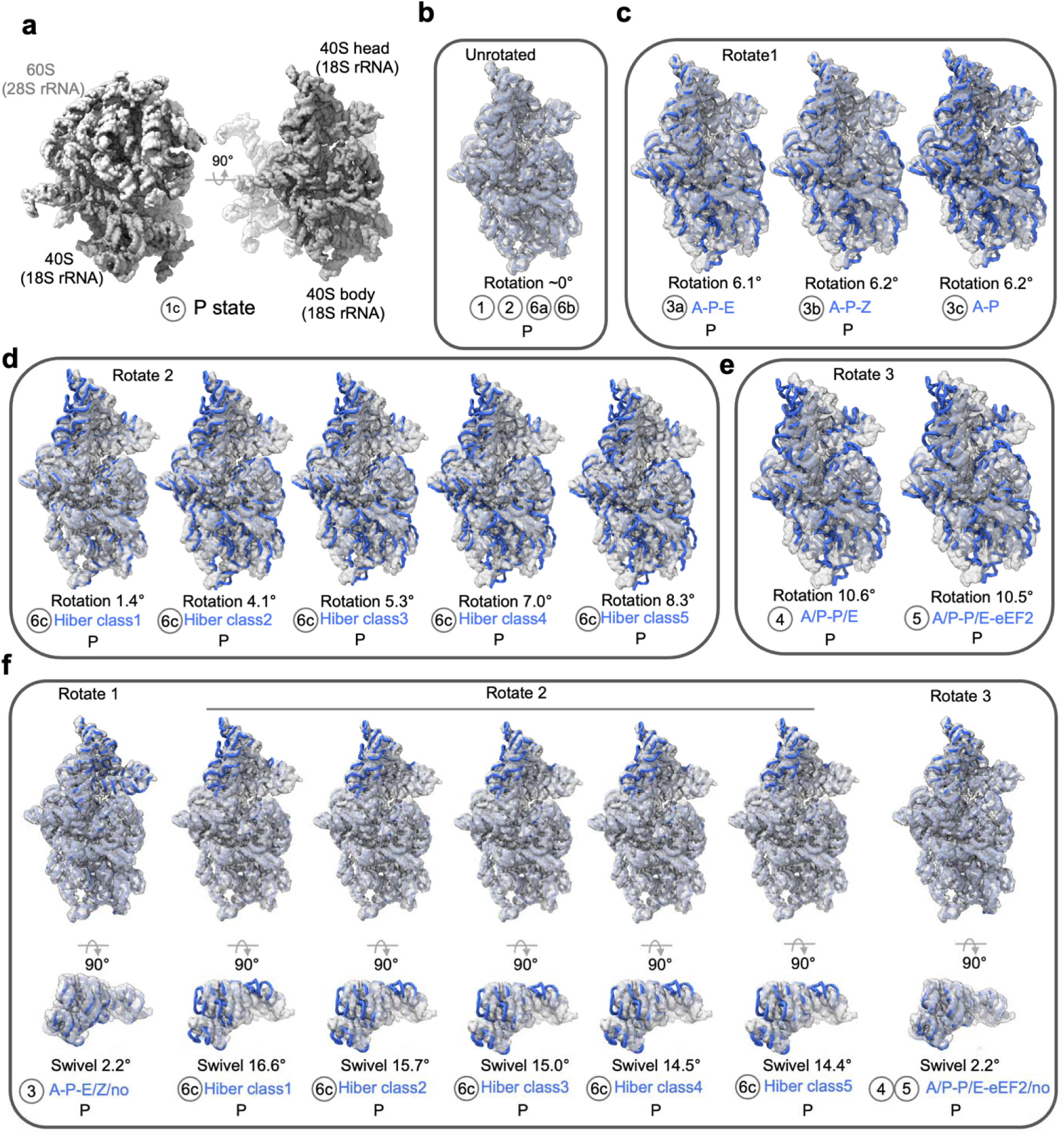
The 40S subunit body domain rotation and head domain swivel for the untreated ribosome states. **a,** The model (28S rRNA and 16S rRNA, surface representation) of the unrotated P state (1c) containing the 60S subunit, 40S subunit body and head domains. **b-e** Comparisons of 40S subunit body domain rotation of different states (ribbon) with unrotated P state (transparent density). **f**. Comparisons of 40S subunit head domain swivel of different states (ribbon) with unrotated P state (transparent density).

**Extended Data Fig. 6:**
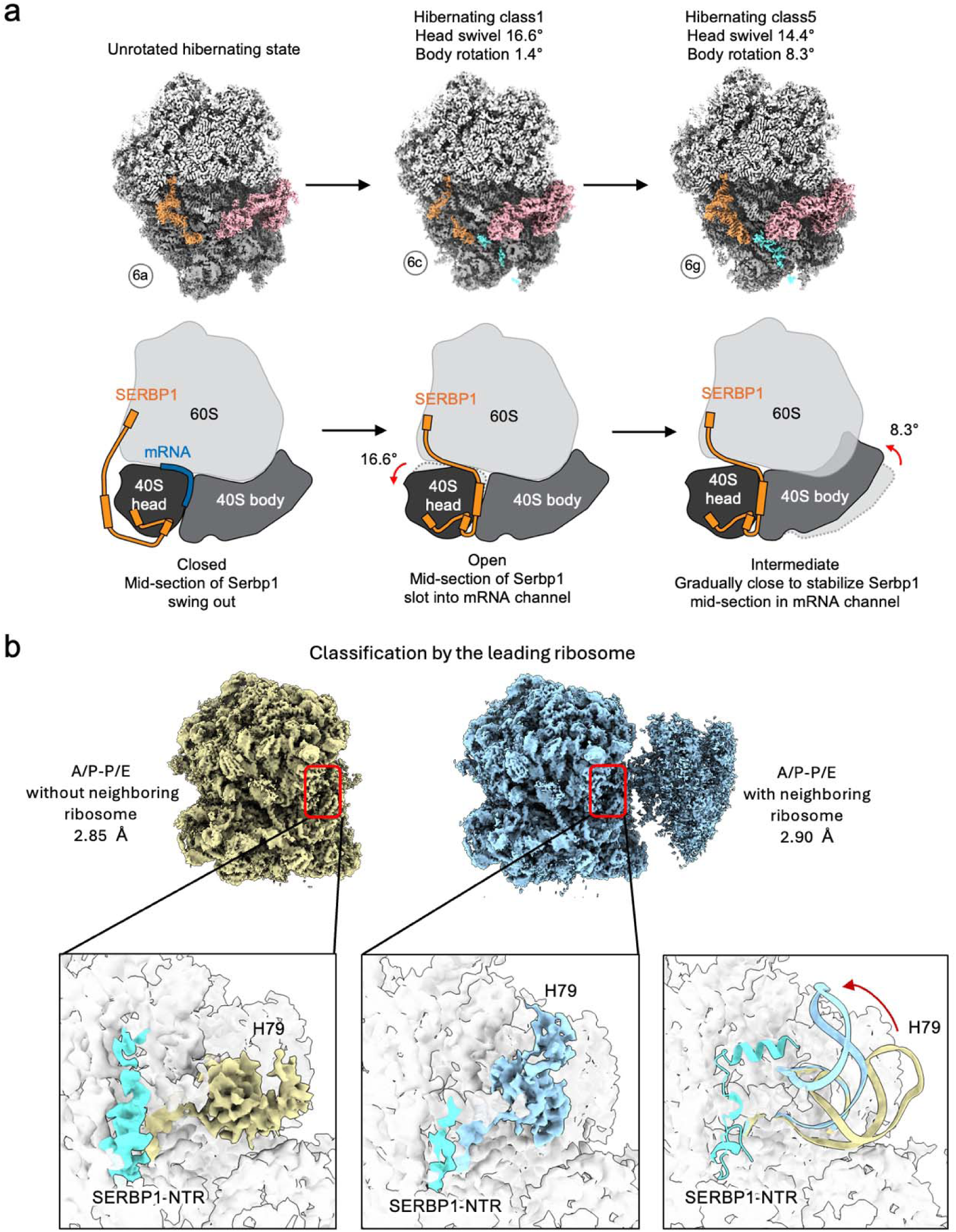
SERBP1 is involved in translation regulation and ribosome collision. **a,** cryo-EM maps (top) and cartoon showing the N- and C-terminal regions (NTR and CTR) of SERBP1 binding to the ribosome with its middle region swinging in and out the mRNA channel of the ribosome. During the hibernating states formation, the 40S subunit head domain is rotated first to allow the mRNA channel opening and SERBP1 mid-section slotting into the channel, then the 40S subunit body domain is rotated to close the channel, which stabilizes the hibernating states. **b**. In A/P-P/E states, the EM density of SERBP1 NTR is weaker (bottom middle) when there is a collision with a neighboring ribosome, upon which H79 of the 28S rRNA near SERBP1 in the leading ribosome have a conformational change.

**Extended Data Fig. 7:**
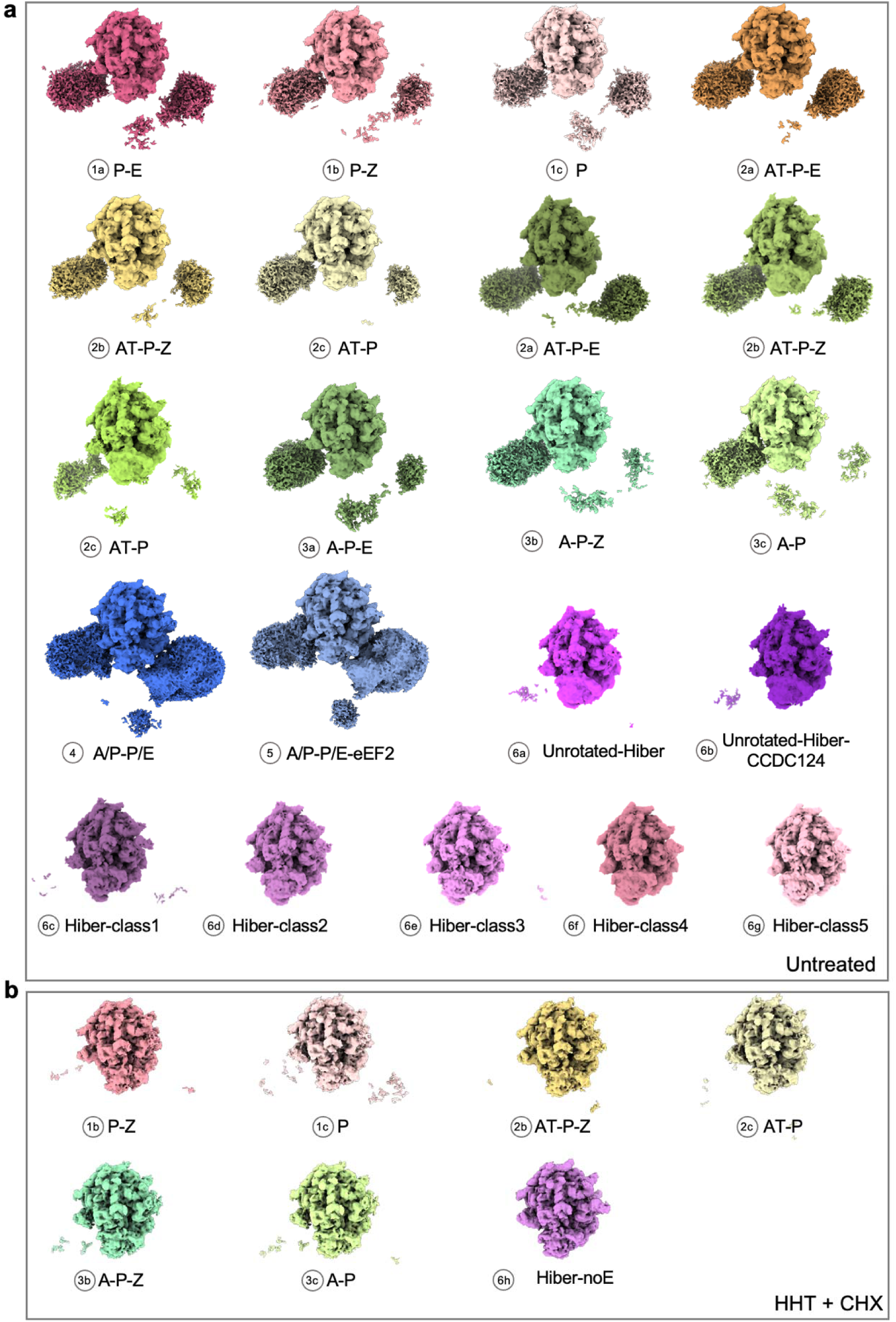
Cryo-EM maps of *in situ* polysomes of different ribosome states. **a,** Densities of the polysome of different states in untreated cells. The particles of different states were re-extracted using a large box size (bin4, 240 pixels) and refined. The maps are displayed using the same threshold. **b,** Same as in a, for HHT+CHX-treated cells.

**Extended Data Fig. 8:**
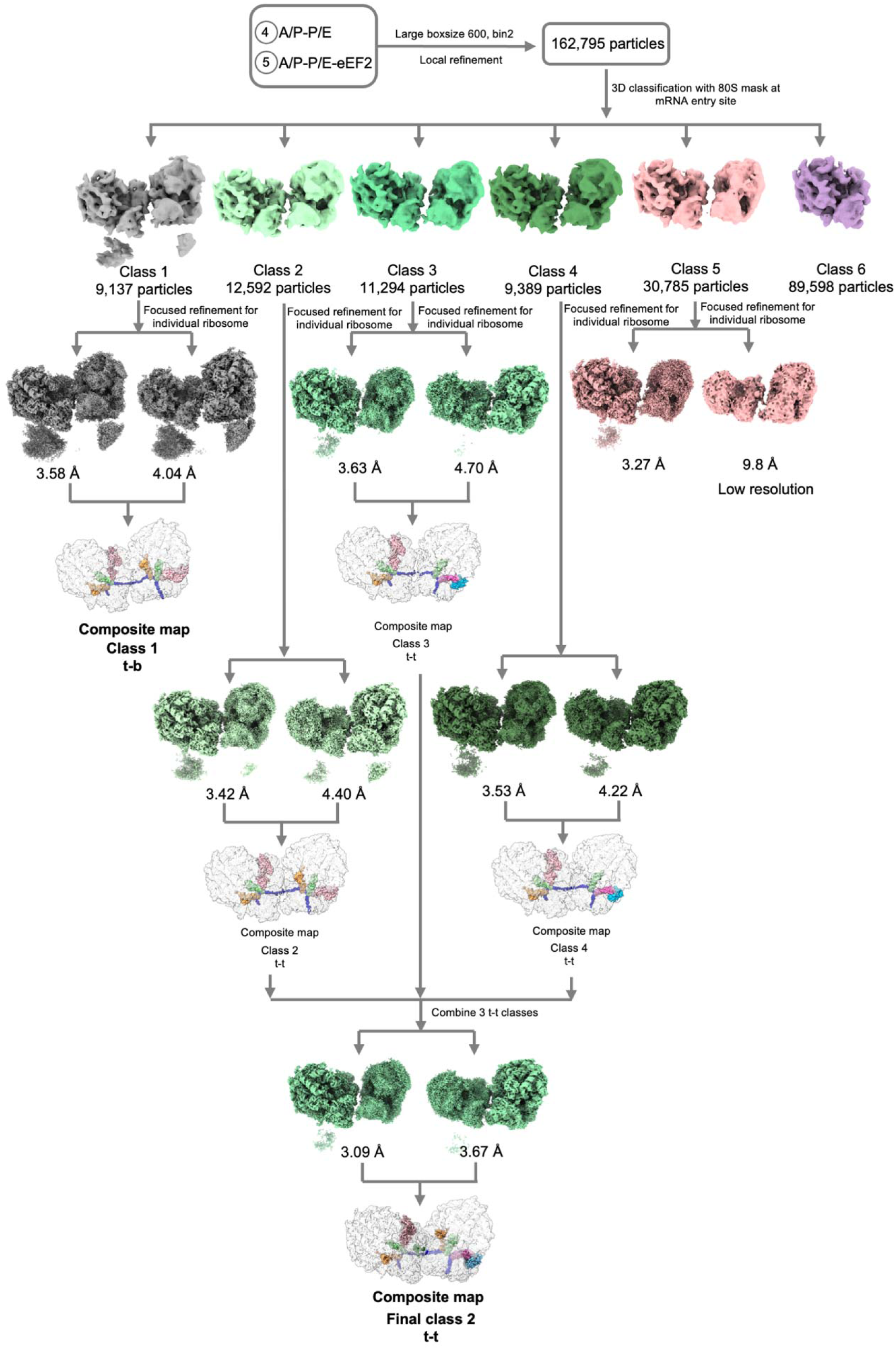
Image processing workflow of the di-ribosome reconstructions.

**Extended Data Fig. 9:**
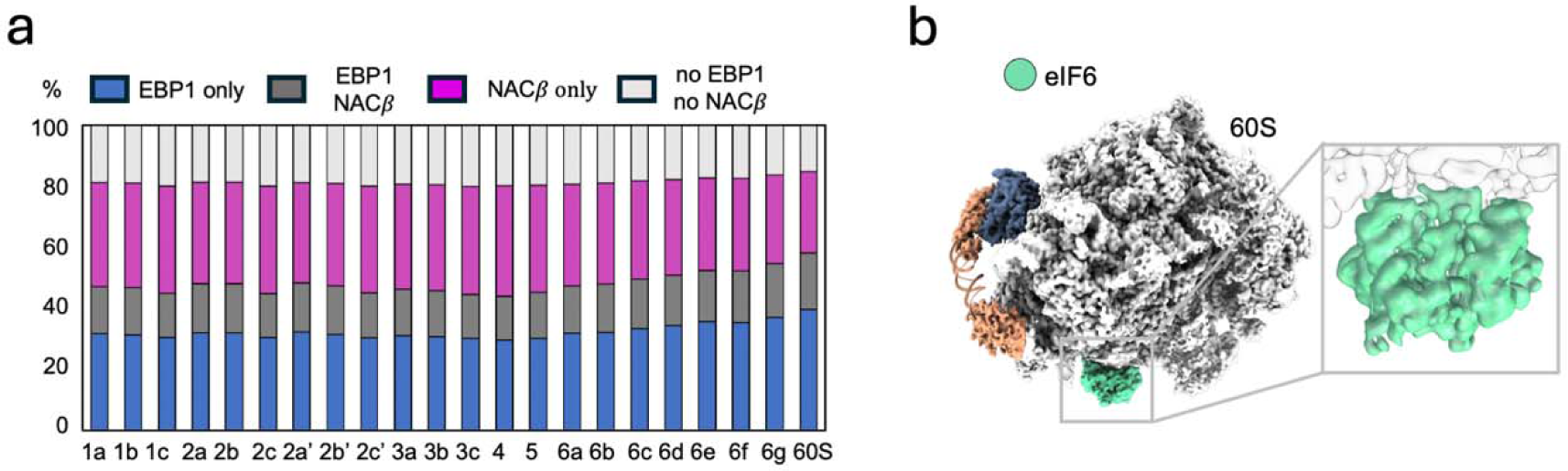
EBP1 and NAC occupancy in different states and eIF6 density in free 60S. a) The EBP1 and NAC occupancies of different 80S conformations and free 60S subunits in the untreated cells. b) The EIF6 (green) and EBP1 densities in free 60S particles in the untreated cells.

**Extended Data Fig. 10:**
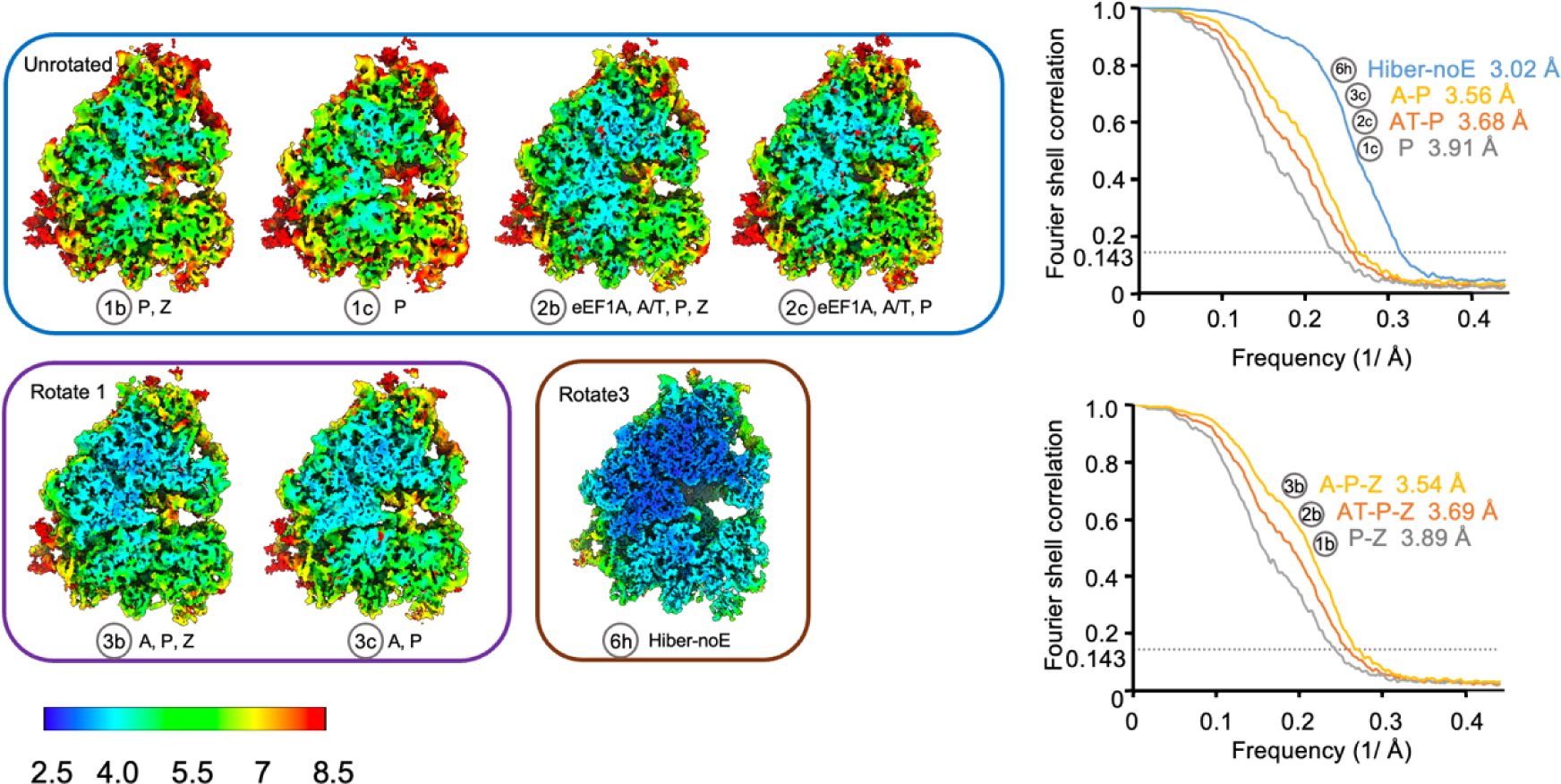
Local resolution maps and FSC of ribosome states of the HHT+CHX-treated cells. FSC curves using the 0.143 criterion.

**Extended Data Fig. 11:**
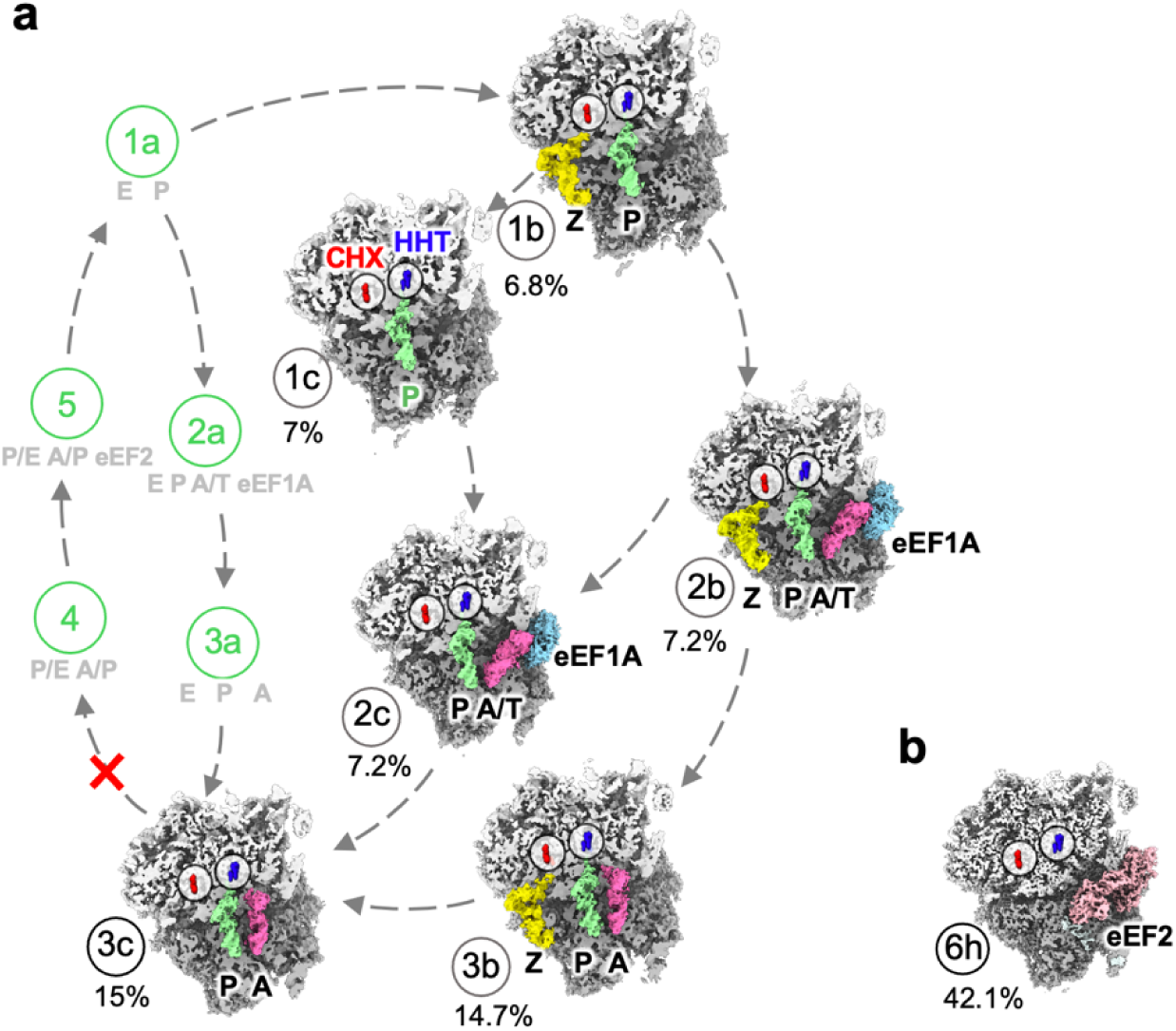
HHT and CHX alter translation dynamics. **a**, The translation states obtained from the HHT+CHX-treated cells, displayed as in Fig. 2a. CHX and HHT densities are highlighted in red and blue respectively. **b**, Only one hibernating state was observed from the CHX+HHT-treated cells.

**Extended Data Table 1-1.**
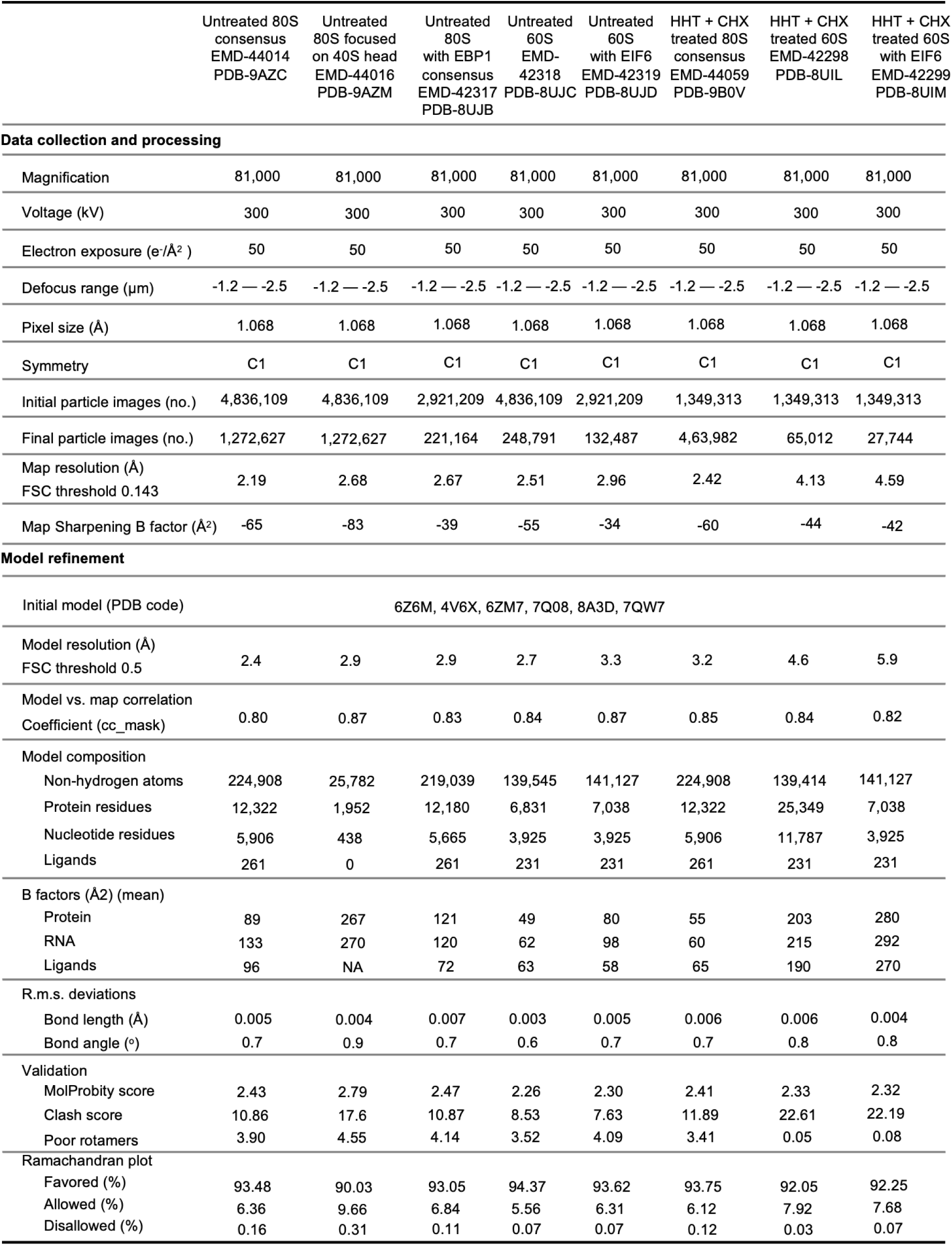
Cryo-EM data collection, refinement and validation statistics of 80S and free 60S consensus structures for untreated and treated datasets.

**Extended Data Table 1-2.**
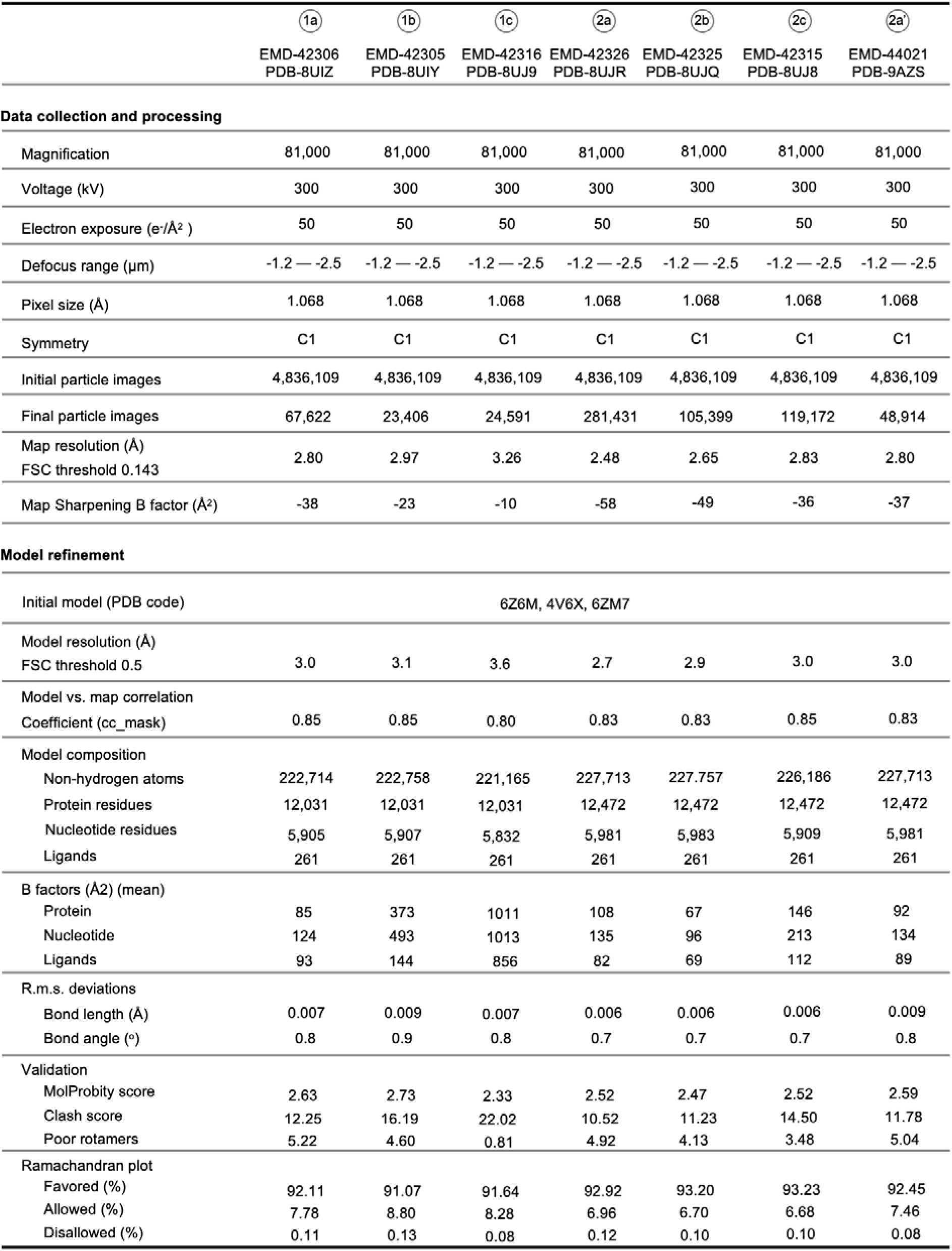
Cryo-EM data collection, refinement and validation statistics of 80S states for untreated dataset (part1).

**Extended Data Table 1-3.** Cryo-EM data collection, refinement and validation statistics of 80S states for untreated dataset (part2).

|  | (2b) | (2c) | (3a) | (3b) | (3c) | (4) | (5) |
| --- | --- | --- | --- | --- | --- | --- | --- |
|  | EMD-44048<br>PDB-9B0O | EMD-44047<br>PDB-9B0N | EMD-42312<br>PDB-8UJ5 | EMD-42311<br>PDB-8UJ4 | EMD-42310<br>PDB-8UJ3 | EMD-44017<br>PDB-9AZN | EMD-44060<br>PDB-9B0W |
| **Data collection and processing** |  |  |  |  |  |  |  |
| Magnification | 81,000 | 81,000 | 81,000 | 81,000 | 81,000 | 81,000 | 81,000 |
| Voltage (kV) | 300 | 300 | 300 | 300 | 300 | 300 | 300 |
| Electron exposure (e-/Å<sup>2</sup>) | 50 | 50 | 50 | 50 | 50 | 50 | 50 |
| Defocus range (µm) | -1.2 — -2.5 | -1.2 — -2.5 | -1.2 — -2.5 | -1.2 — -2.5 | -1.2 — -2.5 | -1.2 — -2.5 | -1.2 — -2.5 |
| Pixel size (Å) | 1.068 | 1.068 | 1.068 | 1.068 | 1.068 | 1.068 | 1.068 |
| Symmetry | C1 | C1 | C1 | C1 | C1 | C1 | C1 |
| Initial particle images | 4,836,109 | 4,836,109 | 4,836,109 | 4,836,109 | 4,836,109 | 4,836,109 | 4,836,109 |
| Final particle images | 50,321 | 60,393 | 86,538 | 64,856 | 66,380 | 36,954 | 130,154 |
| Map resolution (Å) |  |  |  |  |  |  |  |
| FSC threshold 0.143 | 2.83 | 2.92 | 2.75 | 2.83 | 2.93 | 2.98 | 2.66 |
| Map Sharpening B factor (Å<sup>2</sup>) | -34 | -27 | -41 | -34 | -27 | -23 | -49 |
| **Model refinement** |  |  |  |  |  |  |  |
| Initial model (PDB code) | 6Z6M, 4V6X, 6ZM7 |  |  |  |  |  |  |
| Model resolution (Å) |  |  |  |  |  |  |  |
| FSC threshold 0.5 | 3.0 | 3.1 | 3.0 | 3.0 | 3.1 | 3.2 | 2.9 |
| Model vs. map correlation |  |  |  |  |  |  |  |
| Coefficient (cc_mask) | 0.85 | 0.84 | 0.80 | 0.84 | 0.83 | 0.85 | 0.81 |
| Model composition |  |  |  |  |  |  |  |
| Non-hydrogen atoms | 227,757 | 226,164 | 224,272 | 224,272 | 222,679 | 223,564 | 230,169 |
| Protein residues | 12,472 | 12,472 | 12,031 | 12,031 | 12,031 | 12,152 | 12998 |
| Nucleotide residues | 5983 | 5908 | 5,978 | 5,978 | 5,903 | 5902 | 5902 |
| Ligands | 261 | 261 | 261 | 261 | 261 | 261 | 261 |
| B factors (Å<sup>2</sup>) (mean) |  |  |  |  |  |  |  |
| Protein | 138 | 202 | 107 | 139 | 169 | 402 | 77 |
| Nucleotide | 199 | 311 | 163 | 189 | 246 | 542 | 119 |
| Ligands | 77 | 127 | 100 | 127 | 158 | 154 | 81 |
| R.m.s. deviations |  |  |  |  |  |  |  |
| Bond length (Å) | 0.006 | 0.005 | 0.007 | 0.006 | 0.008 | 0.007 | 0.004 |
| Bond angle (°) | 0.7 | 0.7 | 0.8 | 0.7 | 0.8 | 0.8 | 0.8 |
| Validation |  |  |  |  |  |  |  |
| MolProbity score | 2.59 | 2.58 | 2.59 | 2.61 | 2.73 | 2.74 | 2.53 |
| Clash score | 13.72 | 13.68 | 14.31 | 15.13 | 19.39 | 18.70 | 14.21 |
| Poor rotamers | 4.34 | 4.36 | 4.07 | 4.16 | 4.17 | 4.36 | 3.79 |
| Ramachandran plot |  |  |  |  |  |  |  |
| Favored (%) | 92.83 | 92.88 | 92.69 | 92.85 | 92.34 | 92.13 | 93.32 |
| Allowed (%) | 7.07 | 7.03 | 7.21 | 7.06 | 7.59 | 7.71 | 6.52 |
| Disallowed (%) | 0.10 | 0.09 | 0.10 | 0.08 | 0.08 | 0.16 | 0.16 |

**Extended Data Table 1-4.** Cryo-EM data collection, refinement and validation statistics of 80S states for untreated dataset (part3).

|  | 6a | 6b | 6c | 6d | 6e | 6f | 6g |
| --- | --- | --- | --- | --- | --- | --- | --- |
|  | EMD-44063<br>PDB-9B11 | EMD-44043<br>PDB-9B0J | EMD-44049<br>PDB-9B0P | EMD-44051<br>PDB-9B0R | EMD-44039<br>PDB-9B0F | EMD-44040<br>PDB-9B0G | EMD-44041<br>PDB-9B0H |
| **Data collection and processing** |  |  |  |  |  |  |  |
| Magnification | 81,000 | 81,000 | 81,000 | 81,000 | 81,000 | 81,000 | 81,000 |
| Voltage (kV) | 300 | 300 | 300 | 300 | 300 | 300 | 300 |
| Electron exposure (e<sup>-</sup>/Å<sup>2</sup>) | 50 | 50 | 50 | 50 | 50 | 50 | 50 |
| Defocus range (μm) | -1.2 — -2.5 | -1.2 — -2.5 | -1.2 — -2.5 | -1.2 — -2.5 | -1.2 — -2.5 | -1.2 — -2.5 | -1.2 — -2.5 |
| Pixel size (Å) | 1.068 | 1.068 | 1.068 | 1.068 | 1.068 | 1.068 | 1.068 |
| Symmetry | C1 | C1 | C1 | C1 | C1 | C1 | C1 |
| Initial particle images | 4,836,109 | 4,836,109 | 4,836,109 | 4,836,109 | 4,836,109 | 4,836,109 | 4,836,109 |
| Final particle images | 103,321 | 86,934 | 54,349 | 59,299 | 54,480 | 219,754 | 123,628 |
| Map resolution (Å)<br>FSC threshold 0.143 | 2.77 | 2.84 | 2.82 | 2.81 | 2.78 | 2.57 | 2.59 |
| Map Sharpening B factor (Å<sup>2</sup>) | -40 | -36 | -34 | -38 | -39 | -56 | -52 |
| **Model refinement** |  |  |  |  |  |  |  |
| Initial model (PDB code) | 6Z6M, 4V6X, 6ZM7 |  |  |  |  |  |  |
| Model resolution (Å)<br>FSC threshold 0.5 | 3.0 | 3.0 | 3.0 | 3.0 | 3.0 | 2.8 | 2.8 |
| Model vs. map correlation<br>Coefficient (cc_mask) | 0.82 | 0.82 | 0.81 | 0.82 | 0.83 | 0.77 | 0.81 |
| Model composition |  |  |  |  |  |  |  |
| Non-hydrogen atoms | 227,743 | 228,356 | 229,857 | 229,857 | 229,857 | 229,857 | 229,857 |
| Protein residues | 12,877 | 12,950 | 13,151 | 13,151 | 13,151 | 13,151 | 13,151 |
| Nucleotide residues | 5,831 | 5,831 | 5,831 | 5,831 | 5,831 | 5,831 | 5,831 |
| Ligands | 261 | 261 | 261 | 261 | 261 | 261 | 261 |
| B factors (Å<sup>2</sup>) (mean) |  |  |  |  |  |  |  |
| Protein | 77 | 86 | 80 | 81 | 81 | 105 | 117 |
| Nucleotide | 113 | 141 | 133 | 134 | 129 | 152 | 167 |
| Ligands | 81 | 90 | 81 | 100 | 90 | 98 | 107 |
| R.m.s. deviations |  |  |  |  |  |  |  |
| Bond length (Å) | 0.007 | 0.006 | 0.004 | 0.006 | 0.006 | 0.003 | 0.005 |
| Bond angle (°) | 0.8 | 0.8 | 0.7 | 0.7 | 0.8 | 0.7 | 0.7 |
| Validation |  |  |  |  |  |  |  |
| MolProbity score | 2.59 | 2.71 | 2.64 | 2.60 | 2.61 | 2.54 | 2.54 |
| Clash score | 14.79 | 14.88 | 13.96 | 14.09 | 13.37 | 13.50 | 13.33 |
| Poor rotamers | 3.89 | 5.33 | 4.91 | 4.43 | 4.69 | 4.19 | 4.12 |
| Ramachandran plot |  |  |  |  |  |  |  |
| Favored (%) | 92.62 | 92.07 | 92.75 | 92.81 | 92.63 | 93.43 | 93.21 |
| Allowed (%) | 7.22 | 7.79 | 7.12 | 7.08 | 7.24 | 6.46 | 6.67 |
| Disallowed (%) | 0.17 | 0.14 | 0.13 | 0.12 | 0.13 | 0.11 | 0.12 |

**Extended Data Table 1-5.** Cryo-EM data collection, refinement and validation statistics of 80S states for HHT + CHX treated dataset.

|  | (1b) | (1c) | (2b) | (2c) | (3b) | (3c) | (6h) |
| --- | --- | --- | --- | --- | --- | --- | --- |
|  | EMD-42339<br>PDB-8UK0 | EMD-42324<br>PDB-8UJP | EMD-42323<br>PDB-8UJO | EMD-42322<br>PDB-8UJN | EMD-42321<br>PDB-8UJK | EMD-42320<br>PDB-8UJJ | EMD-42351<br>PDB-8UKB |
| **Data collection and processing** |  |  |  |  |  |  |  |
| Magnification | 81,000 | 81,000 | 81,000 | 81,000 | 81,000 | 81,000 | 81,000 |
| Voltage (kV) | 300 | 300 | 300 | 300 | 300 | 300 | 300 |
| Electron exposure (e-/Å<sup>2</sup>) | 50 | 50 | 50 | 50 | 50 | 50 | 50 |
| Defocus range (μm) | -1.2 — -2.5 | -1.2 — -2.5 | -1.2 — -2.5 | -1.2 — -2.5 | -1.2 — -2.5 | -1.2 — -2.5 | -1.2 — -2.5 |
| Pixel size (Å) | 1.068 | 1.068 | 1.068 | 1.068 | 1.068 | 1.068 | 1.068 |
| Symmetry | C1 | C1 | C1 | C1 | C1 | C1 | C1 |
| Initial particle images | 1,053,496 | 1,053,496 | 1,053,496 | 1,053,496 | 1,053,496 | 1,053,496 | 1,053,496 |
| Final particle images | 7,312 | 7,052 | 9,091 | 8,604 | 15,150 | 14,418 | 52,735 |
| Map resolution (Å) |  |  |  |  |  |  |  |
| FSC threshold 0.143 | 3.89 | 3.91 | 3.69 | 3.68 | 3.54 | 3.56 | 3.05 |
| Map Sharpening B factor (Å<sup>2</sup>) | 0 | 0 | 0 | 0 | -5 | -3 | -30 |
| **Model refinement** |  |  |  |  |  |  |  |
| Initial model (PDB code) | 6Z6M, 4V6X, 6ZM7, 7Q08, 8A3D |  |  |  |  |  |  |
| Model resolution (Å) |  |  |  |  |  |  |  |
| FSC threshold 0.5 | 4.4 | 4.4 | 4.1 | 4.1 | 4.0 | 4.0 | 3.3 |
| Model vs. map correlation |  |  |  |  |  |  |  |
| Coefficient (cc_mask) | 0.84 | 0.85 | 0.85 | 0.85 | 0.83 | 0.85 | 0.85 |
| Model composition |  |  |  |  |  |  |  |
| Non-hydrogen atoms | 222,758 | 221,165 | 227,779 | 226,186 | 224,272 | 222,679 | 226,590 |
| Protein residues | 12,031 | 12,031 | 12,472 | 12,472 | 12,031 | 12,031 | 12,932 |
| Nucleotide residue | 5,907 | 5,832 | 5,984 | 5,909 | 5,978 | 5,903 | 5,756 |
| Ligands | 261 | 261 | 261 | 261 | 261 | 261 | 261 |
| B factors (Å<sup>2</sup>) (mean) |  |  |  |  |  |  |  |
| Protein | 208 | 212 | 180 | 187 | 132 | 177 | 142 |
| Nucleotide | 201 | 200 | 177 | 179 | 164 | 171 | 145 |
| Ligands | 148 | 145 | 126 | 127 | 115 | 121 | 87 |
| R.m.s. deviations |  |  |  |  |  |  |  |
| Bond length (Å) | 0.006 | 0.008 | 0.005 | 0.006 | 0.007 | 0.007 | 0.007 |
| Bond angle (°) | 0.9 | 1.0 | 0.8 | 0.9 | 0.9 | 0.9 | 0.8 |
| Validation |  |  |  |  |  |  |  |
| MolProbity score | 2.20 | 2.43 | 2.20 | 2.29 | 2.24 | 2.25 | 2.08 |
| Clash score | 16.52 | 25.70 | 16.08 | 18.61 | 16.96 | 17.49 | 12.13 |
| Poor rotamers | 0.69 | 1.02 | 0.87 | 0.88 | 0.86 | 0.71 | 0.76 |
| Ramachandran plot |  |  |  |  |  |  |  |
| Favored (%) | 92.22 | 90.85 | 91.93 | 90.08 | 91.37 | 91.25 | 92.01 |
| Allowed (%) | 7.70 | 9.06 | 8.08 | 9.00 | 8.51 | 8.65 | 7.86 |
| Disallowed (%) | 0.08 | 0.08 | 0.12 | 0.11 | 0.12 | 0.10 | 0.13 |

**Extended Data Table 1-6.** Cryo-EM data collection, refinement and validation statistics of disome structures for untreated dataset.

|  | T-t disome Composite<br>EMD-44052<br>PDB-9B0S<br>Focused leading<br>EMD-44057<br>Focused trailing<br>EMD-44056 | T-b disome composite<br>EMD-44050<br>PDB-9B0Q<br>Focused leading<br>EMD-44054<br>Focused trailing<br>EMD-44053 |
| --- | --- | --- |
| **Data collection and processing** |  |  |
| Magnification | 81,000 | 81,000 |
| Voltage (kV) | 300 | 300 |
| Electron exposure (e<sup>-</sup>/Å<sup>2</sup>) | 50 | 50 |
| Defocus range (μm) | -1.2 — -2.5 | -1.2 — -2.5 |
| Pixel size (Å) | 1.068 | 1.068 |
| Symmetry | C1 | C1 |
| Initial particle images | 4,836,109 | 4,836,109 |
| Final particle images | 33,275 | 9,137 |
| Map resolution (Å) | 3.4 | 3.8 |
| FSC threshold 0.143 |  |  |
| Map Sharpening B factor (Å<sup>2</sup>) | -19 | -4 |
| **Model refinement** |  |  |
| Initial model (PDB code) | 6Z6M, 4V6X, 6ZM7 |  |
| Model resolution (Å) | 3.6 | 4.1 |
| FSC threshold 0.5 |  |  |
| Model vs. map correlation |  |  |
| Coefficient (cc_mask) | 0.77 | 0.80 |
| Model composition |  |  |
| Non-hydrogen atoms | 458,132 | 459,641 |
| Protein residues | 25,356 | 25,993 |
| Nucleotide residue | 11,938 | 11,740 |
| Ligands | 522 | 522 |
| B factors (Å<sup>2</sup>) (mean) |  |  |
| Protein | 490 | 1032 |
| Nucleotide | 728 | 1000 |
| Ligands | 260 | 423 |
| R.m.s. deviations |  |  |
| Bond length (Å) | 0.005 | 0.006 |
| Bond angle (°) | 0.8 | 0.8 |
| Validation |  |  |
| MolProbity score | 2.28 | 2.44 |
| Clash score | 20.39 | 28.00 |
| Poor rotamers | 0.52 | 0.47 |
| Ramachandran plot |  |  |
| Favored (%) | 92.19 | 91.52 |
| Allowed (%) | 7.70 | 8.34 |
| Disallowed (%) | 0.12 | 0.14 |

**Extended Data Table 2.** Polyamines identified in the ribosome.

| Number | This Study | 8GLP | 8A3D | Interactions |
| --- | --- | --- | --- | --- |
| L5/5313 | SPM | SPD | SPM | 28S A1527, 28S U1528, 28S G3918, 28S A4186, 28S G4187, L15 Y81 |
| L5/5314 | SPD | SPD | Mg | 28S A39, 28S G40, 28S C41, 28S G91, 28S C92 28S G93, 28S U4340, 28S C4341, L27a H39 |
| L5/5315 | SPM | 2 Mg + K | 2 Mg + K | 28S A44, 28S U45, 28S A47, 28S G48, 28S G90, 28S U291, 28S G293, 28S G294 |
| L5/5316 | SPM | PUT | None | 28S U1735, 28S A1736, 28S A1787, 28S A1788, 28S U1790, 28S U1791, L21 Y13, L21 S16 |
| L5/5317 | SPM | PUT + Mg + K | K | 28S C1678, 28S G1680, 28S G1854, 28S U4194, 28S C4204, 28S 6MZ4220, 28S C4221 |
| L5/5318 | SPD | PUT + Mg | Mg | 28S G1587, 28S G1604, 28S G1605, 28S A3642, 28S A3643 |
| L5/5319 | SPD | SPD | Mg | 28S A64, 28S A328, 28S A329, 28S G330, 28S G331, 28S C332, L15 R169 |
| L5/5320 | SPD | SPD | K | 28S G1581, 28S PSU1582, 28S A1583, 28S G1584, 28S C1585, 28S C1607, 28S G1608, 28S U1609 |
| L5/5321 | SPD | 2 Mg | Mg | 28S G1869, 28S A1939, 28S G3874, 28S G3875, 28S A3877 |
| L5/5322 | SPD | SPD | None | 5.8S G11, 5.8S A12, 5.8S U13, 5.8S C138, 5.8S G139, 5.8S C140 |
| L5/5323 | SPD | SPD | None | 28S A435, 28S C436, 28S C1305, 28S C1306, 28C A1307, 5.8S U7 |
| S2/1930 | SPM | SPD + 2 Mg | N/A | 18S A671, 18S A672, 18S G673, 18S C674, 18S U675, 18S C676, 18S G677, 18S C1026, 18S C1085, 18S U1088, 18S G1089 |
| S2/1931 | SPD | Mg | N/A | 18S PSU105, 18S C106 18S A348, 18S A349, 18S C350, 18S G351, 18S G431, 18S G432 |
| S2/1932 | SPD | Mg | N/A | 18S C1529, 18S A1651, 18S G1652, 18S U1653 |

